# Obesity impairs cognitive function via metabolic syndrome and cerebrovascular disease: an SEM analysis in 15,000 adults from the UK Biobank

**DOI:** 10.1101/2020.06.26.174086

**Authors:** Filip Morys, Mahsa Dadar, Alain Dagher

## Abstract

Chronic obesity is associated with several complications, including cognitive impairment and dementia. However, we have piecemeal knowledge of the mechanisms linking obesity to central nervous system damage. Adiposity leads to the metabolic syndrome, consisting of inflammation, hypertension, dyslipidemia and insulin resistance. In turn, these metabolic abnormalities cause cerebrovascular dysfunction, which may cause white and grey matter tissue loss and consequent cognitive impairment. While there have been several neuroimaging studies linking adiposity to changes in brain morphometry, a comprehensive investigation of the relationship has so far not been done. Here we use structural equation modelling applied to over 15,000 individuals from the UK Biobank to identify the causal chain that links adiposity to cognitive dysfunction. We found that body mass index and waist-to-hip ratio were positively related to higher plasma C-reactive protein, dyslipidemia, occurrence of hypertension and diabetes, all of which were in turn related to cerebrovascular disease as measured by volume of white matter hyperintensities on magnetic resonance imaging. White mater hyperintensities were associated with lower cortical thickness and volume and higher subcortical volumes, which were associated with cognitive deficits on tests of visuospatial memory, fluid intelligence, and working memory among others. In follow-up analyses we found that inflammation, hypertension and diabetes mediated 20% of the relationship between obesity and cerebrovascular disease and that cerebrovascular disease mediated a significant proportion of the relationship between obesity and cortical thickness and volume. We also showed that volume of white matter hyperintensities was related to decreased fractional anisotropy and increased mean diffusivity in the majority of white matter tracts, pointing to white matter dysconnectivity as a major cause of impaired cognition. Our results have clinical implications, supporting a role for the management of adiposity in the prevention of late-life dementia and cognitive decline.

## 1 Introduction

Obesity is a significant human and financial burden (Apovian, 2016; Hammond and Levine, 2010). Among its many adverse health outcomes, chronic obesity in midlife is associated with subsequent cognitive impairment, and is a risk factor for vascular dementia and Alzheimer’s Disease (Whitmer et al., 2007).

Obesity is defined as the excessive accumulation of adipose tissue. Its link with cognitive dysfunction has been attributed to the metabolic syndrome, which consists of inflammation, hypertension, dyslipidemia, and insulin resistance (Yaffe et al., 2004). Adipose tissue releases proinflammatory cytokines, such as interleukin-6 (IL-6), and inflammation-related proteins, such as C-reactive protein (CRP), which, in obesity, lead to low-grade systemic inflammation (Monteiro and Azevedo, 2010; Sproston and Ashworth, 2018; Visser et al., 2001). Endocrine activities of adipose tissue also increase the risk of diabetes, hypertension and dyslipidemia (Cusi, 2010; Hwang et al., 2016; Zhou and Qin, 2012). These comorbidities are causally related to cerebrovascular dysfunction (Bowman et al., 2012; Dufouil et al., 2001; Hsuchou et al., 2012a; Lampe et al., 2018; Tamura and Araki, 2015; Wardlaw et al., 2015, 2017). Vascular damage in white matter may lead to demyelination, loss of oligodendrocytes, and gliosis (Aribisala et al., 2013; Bailey et al., 2012; Gouw et al., 2011; Hsuchou et al., 2012a; Lambert et al., 2015; Pantoni, 2010; Wardlaw et al., 2017).

In magnetic resonance imaging (MRI), small vessel cerebrovascular disease manifests as white matter hyperintensities (WMH) on T2-weighted imaging and changes in white matter integrity measured by diffusion-weighted imaging (Hakim, 2019; Walker et al., 2017; Wardlaw et al., 2015). White matter disruption has been associated with grey matter atrophy in cross-sectional studies (Lambert et al., 2015; Pareek et al., 2018; Tuladhar et al., 2015). There is evidence that WMH precede cortical thinning from a longitudinal study in Parkinson’s disease (Dadar et al., 2018). Obesity was previously associated with cortical and subcortical functional and volumetric alterations (Beyer et al., 2019a; García-García et al., 2018, 2019; García-García et al., 2019; Horstmann et al., 2011, 2015; Neseliler et al., 2018; Pannacciulli et al., 2006; Rapuano et al., 2017; Vainik et al., 2018; Volkow et al., 2008). These changes, together with white matter microstructure alterations, lead to poorer cognitive performance (Beyer et al., 2019a; Kharabian Masouleh et al., 2016; Samara et al., 2020; Zhang et al., 2018). Cognitive domains affected are executive function, verbal memory, processing speed (Beyer et al., 2019a; Kharabian Masouleh et al., 2016; Samara et al., 2020; Zhang et al., 2018), fluid intelligence (Spyridaki et al., 2014), and working memory (Alarcón et al., 2016). In sum, adiposity seems to affect white and grey matter integrity, and hence cognitive function, possibly via low-grade systemic inflammation, hypertension, diabetes and dyslipidemia (Lampe et al., 2018; Miller and Spencer, 2014; Nguyen et al., 2014; Zhang et al., 2018).

While separate links between obesity and white matter damage, obesity and grey matter atrophy, and obesity and poor cognitive performance are established, no study has investigated the proposed pathway in full. In this study, we set out to investigate this mechanism in one model with the following hypotheses: 1) obesity measures, such as body mass index (BMI) and waist-to-hip ratio (WHR), are positively related to inflammation, hypertension, diabetes, triglycerides levels, and negatively to high density lipoprotein levels (dyslipidemia), which in turn affect WMH volume; 2) WMH load affects grey matter morphometry; 3) all these variables may be independently associated with cognitive performance. To test this, we first utilised structural equation modeling (SEM), creating a model in line with our hypotheses. Next, we followed up on all significant associations using mediation analyses to better understand precise relationships between single variables, and to test whether obesity is a causal factor for these associations. Finally, we investigated how WMH load is related to white matter microstructure measures – fractional anisotropy and mean diffusivity – to pinpoint mechanisms in which white matter damage potentially contributes to cognitive deficits and grey matter changes.

We used both BMI and WHR as obesity measures as they differentially predict obesity-related outcomes (Debette et al., 2010; Dobbelsteyn et al., 2001; Noble, 2001).

Here, we used data from the UK Biobank cohort study and available imaging derived phenotypes (Alfaro-Almagro et al., 2018). The large sample size (∼15 000 participants) enabled us to generate a comprehensive SEM with latent factors combining measures of adiposity, metabolic syndrome, brain abnormalities, and cognitive dysfunction.

## 2 Materials and methods

### 2.1 Sample characteristic

Here, we used the UK Biobank dataset – a large scale study with extensive phenotyping and brain imaging data (Miller et al., 2016; Sudlow et al., 2015). This study was performed under UK Biobank application ID 35605. The number of participants with available imaging data was 21,333. Prior to all analyses, we excluded individuals with underweight (BMI<18.5kg/m^2^), bipolar disorder, personality disorder, schizophrenia, mania, bulimia nervosa, anorexia nervosa, ADD/ADHD, panic attacks and neurological illnesses (n=4784, Table S1). UK Biobank field IDs used in this study can be found in Table S2. Presence of diabetes and hypertension in participants was determined based on previous diagnoses. All participants signed informed consents prior to participating in the study, which was approved by the North-West Multi-centre Research Ethics Committee. All UK Biobank actions are overseen by the UK Biobank Ethics Advisory Committee.

### 2.2 Obesity measures

We investigated body mass index and waist-to-hip ratio as measures of adiposity. BMI data were provided directly by the UK Biobank and were calculated as the ratio of weight to height squared. We calculated WHR by dividing waist circumference by hip circumference. Obesity measures were collected at the time of brain imaging.

### 2.3 Imaging derived phenotypes

For brain analyses we used imaging derived phenotypes available from UK Biobank. Imaging pipeline details can be found in Alfaro-Almagro et al. 2018. For cortical grey matter morphometry we used cortical volume and thickness as derived by FreeSurfer (Fischl, 2012) for each parcel of the Desikan-Killiany-Tourville (DKT) atlas (Klein and Tourville, 2012). Subcortical volumes were derived using FMRIB’s Integrated Registration and Segmentation Tool (FIRST; Patenaude et al. 2011; Jenkinson et al. 2012). Here, we investigated volumes of the following subcortical structures: the thalamus, caudate nucleus, putamen, pallidum, hippocampus, amygdala and nucleus accumbens. Volume of WMH was calculated using Brain Intensity AbNormality Classification Algorithm (BIANCA; Griffanti et al. 2016). FA and MD values for each of the white matter tracts were obtained using DTIFIT within FMRIB’s Software Library (FSL; Jenkinson et al. 2012). Both are measures of tissue integrity. FA is a measure of directionality of diffusion of a water molecule within white matter tracts. Low FA is thought to indicate reduced integrity of white matter fibers and/or lower myelination (Alba-Ferrara and de Erausquin, 2013; Alexander et al., 2007; Basser, 1995; Sen and Basser, 2005; Song et al., 2002). MD is a measure of the magnitude of diffusivity of a water molecule in all directions (Madden et al., 2012; Sen and Basser, 2005; Soares et al., 2013). Increased values of MD indicate higher diffusivity of water molecules and might be related to edema, blood brain barrier damage and brain tissue microstructure disruptions (Alexander et al., 2007; Wardlaw et al., 2017).

### 2.4 Blood markers

We used three blood markers of metabolic syndrome: serum C-reactive protein (CRP), a marker of low-grade systemic inflammation (Sproston and Ashworth, 2018) and serum triglyceride and high density lipoprotein (HDL) levels as indicators of dyslipidemia. Detailed description of blood sampling and procedures can be found in Elliott et al. 2008. Blood samples were collected at the initial assessment visit, while other measures used in this study were collected during the first imaging visit, which took place on average 8 years later. However, since our question of interest is how obesity is related to CRP and dyslipidemia and since obesity measures between the two timepoints are highly correlated (BMI: r=0.897, p<0.0001; WHR: 0.791, p<0.0001), we decided to use the CRP, triglycerides and HDL levels from the earlier timepoint. High correlation of obesity measures between the two timepoints indicates that inflammation and dyslipidemia marker values between the two timepoints would also be highly correlated, assuming that they are related to obesity.

### 2.5 Cognitive measures

In UK Biobank, cognitive tests were administered via a touchscreen interface. The tests lasted approximately 15 minutes (Cullen et al., 2015) and are described in detail online at http://biobank.ctsu.ox.ac.uk/crystal/label.cgi?id=100026. Overall, despite the cognitive tests being non-standard, they correspond well with their standardized and well-validated counterparts and exhibit good test-retest reliability (Fawns-Ritchie and Deary, 2019). We investigated the following 6 cognitive abilities: 1) working memory, assessed by digit span task; 2) fluid intelligence, assessed by a set of reasoning tasks; 3) executive functions, measured by the tower rearranging test; 4) prospective memory, assessed by a number of times an intention was forgotten on a prospective memory task; 5) visuospatial memory, assessed by a pairs matching task (here the measurement indicated the number of incorrect pair matchings); 6) reaction time, a measure of processing speed. Some of the cognitive measures were not administered to all participants, hence sample sizes might be smaller (see sections 2.6.1 and 2.6.2.1). Because tests of visuospatial memory and reaction time were administered more than once, we used an average value of all trials. Note that for prospective memory, visuospatial memory and reaction time, higher scores indicate worse performance.

### 2.6 Data analysis

#### 2.6.1 Structural equation model

We used an SEM to test the relation between obesity, metabolic syndrome, and brain and cognitive dysfunction. We prepared the data for the analysis in the following ways: First, volumetric brain measures – volume of WMH and cortical and subcortical grey matter – were corrected for head size. For software consistency, the scaling factor for WMH and subcortical volumes was derived from FSL, while for cortical volumes we used the measure of estimated total intracranial volume from FreeSurfer. Second, BMI, WMH, CRP, HDL and triglycerides levels were log-transformed and all variables were z-scored prior to analysis. Further, we pairwise excluded outliers as defined by datapoints being 2.2 interquartile range below 1^st^ or above 3^rd^ quartile (Tukey 1977; Hoaglin et al. 1986; Hoaglin and Iglewicz 1987), which resulted in a dataset containing 16,438 participants (Table S3) with missing values (which are allowed in the SEM strategy described below). Sample size for each variable used in the SEM can be found in Table S4.

We used the lavaan package in R (version 0.6-5; Rosseel 2012) to define an SEM consisting of 8 latent variables: obesity (consisting of BMI and WHR), the four metabolic syndrome markers, inflammation (serum CRP), dyslipidemia (serum triglyceride and serum HDL), hypertension and diabetes, cerebrovascular disease (WMH), cortical thickness (thickness for all DKT parcels), cortical volume (volume for all DKT parcels), subcortical volume (volume for all measured subcortical parcels), and cognition (consisting of 6 cognitive measures described in section 2.5). A representation of this model can be found in Figure 1. In brief, in our model obesity affected inflammation, dyslipidemia, hypertension, and diabetes, which in turn affected cerebrovascular disease. This led to changes in cortical volume, thickness and subcortical volume. Finally, all measures affected cognition. In each of the regressions we controlled for age and sex. Education level was used as a covariate in all regressions including cognition as an outcome. Because the model contained categorical variables, it was estimated using robust diagonally weighted least squares estimation with pairwise missing values exclusions. SEM p-values for each parameter were corrected for multiple comparisons using an adjusted Bonferroni correction (Smith and Cribbie, 2013).

**Figure 1.**
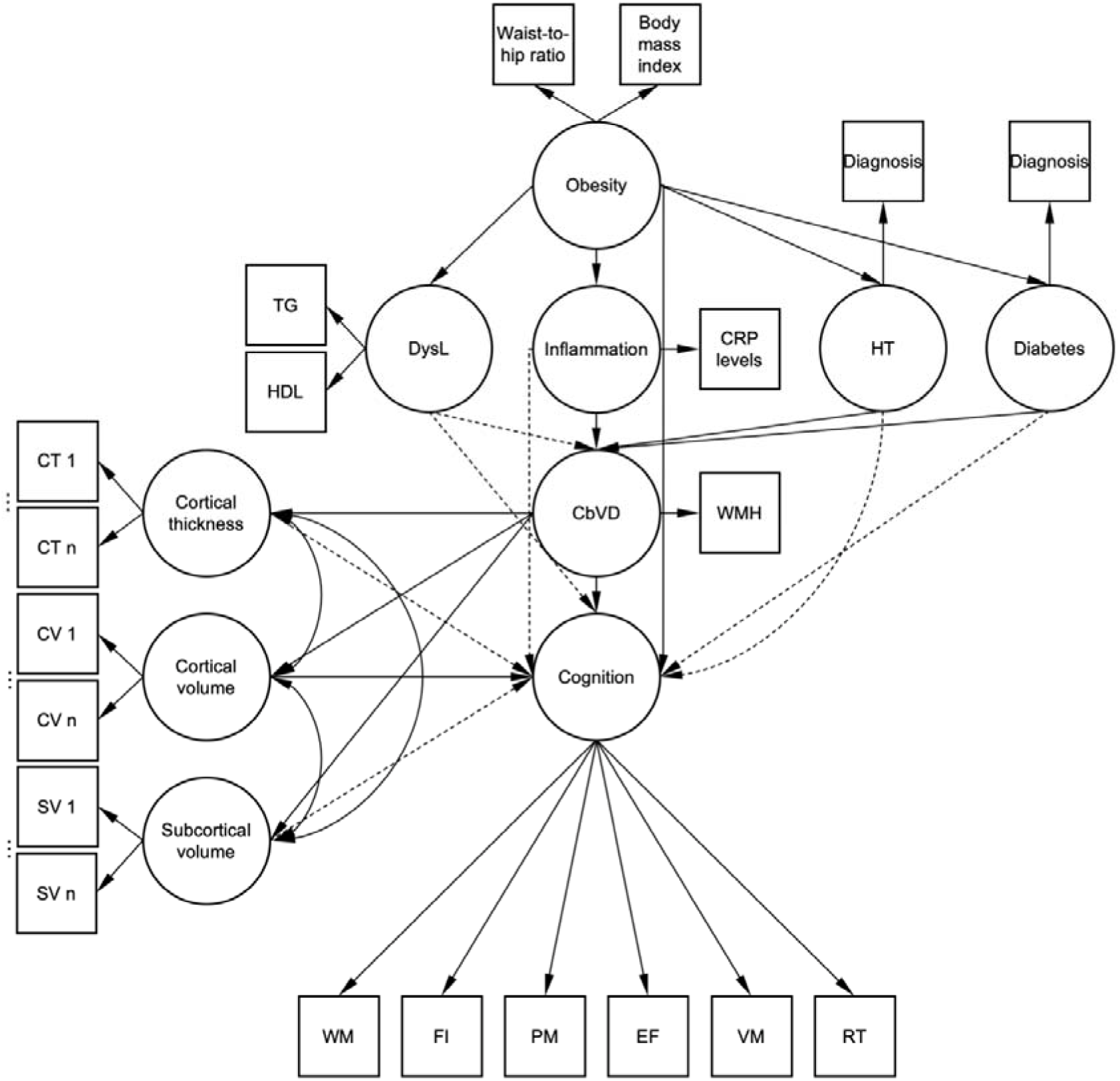
Schematic representation of the structural equation model. Dashed lines represent non-significant associations. Note that not all measured variables contributing to cortical thickness, cortical volume and subcortical volume are depicted in this graph. Age and sex were added to each of the associations between variables as covariates of no interest and are not depicted here. Education was added in each regression with cognition as outcome variable and is not depicted here. DysL – dyslipidemia, TG – triglycerides, HDL – high density lipoprotein, HT – hypertension, CRP – C-reactive protein, CbVD – cerebrovascular disease, WMH – white matter hyperintensities, CT – cortical thickness, CV – cortical volume, SV – subcortical volume, WM – working memory, FI – fluid intelligence, PM – prospective memory, EF – executive function, VM – visuospatial memory, RT – reaction time.

#### 2.6.2 Post-SEM analyses

Having established a relationship between obesity, inflammation, dyslipidemia, brain measures and cognition, we also analysed the variables separately, instead of pooling them into latent variables. This allowed us to investigate different aspects of obesity – BMI and WHR – as well as separate brain structures and parcels, and cognitive measures to investigate associations identified in the SEM in more depth.

We first investigated general associations between obesity measures and grey matter and cognition. We then used mediation analyses to test whether BMI and WHR exert effects on cerebrovascular disease, cognition and grey matter directly or via other mediators and what the patterns of these associations are. Here, we only investigated the relationships that were significant in the SEM.

From the sample described in section 2.1 we excluded BMI, WHR and WMH outliers listwise from analysis (2.2 interquartile range below/above 1^st^ and 3^rd^ quartile, respectively; Tukey 1977; Hoaglin et al. 1986; Hoaglin and Iglewicz 1987). The resulting sample consisted of 15,262 individuals (Table S3). Next, we excluded outliers from our measures of interest. This was done separately for measures of cortical thickness and cortical and subcortical grey matter volumes. However, for the sake of sample size consistency within each of those modalities, any outlier identified in one cortical parcel/subcortical structure was removed in all other parcels/structures. For cognitive measures, due to sample size differences, we decided to exclude outliers separately for each of the measures. Next, volumetric brain measures were corrected for total brain volume. Finally, we log-transformed WMH volume, BMI and CRP values to achieve a normal distribution of these variables. Prior to all analyses, all numerical variables were z-scored. In every analysis step we corrected for age and sex. Initially, we also included age^2^, however it did not significantly alter the results and we decided to exclude it from analyses. All analyses were performed in R (v. 3.6.0). Figures were prepared in MATLAB R2019b and using python (v. 3.7.4) with the nilearn (v. 0.5.1) library (Abraham et al., 2014). Scripts used in the study can be found at https://github.com/FilipMorys/WMHOB.

##### 2.6.2.1 Obesity associations with grey matter morphometry and cognition

We used a general linear model to assess the associations between obesity measures – BMI and WHR – and grey matter morphometry, but also obesity measures and cognition. P-values were corrected for multiple comparisons using FDR Benjamini-Hochberg correction and were considered significant at α=0.05 (Benjamini and Hochberg, 1995). The correction was applied separately for each grey matter morphometry modality, and collectively for all cognitive measures.

Sample sizes for these analyses and the following mediation analyses using the same measures of interest were: 1) cortical thickness: 14,457; 2) cortical volume: 14,117; 3) subcortical volume: 13,945; 4) working memory: 6,720; 5) fluid intelligence: 13,739; 6) executive function: 5,499; 7) prospective memory: 14,112; 8) visuospatial memory: 13,761; 9) reaction time: 13,880.

##### 2.6.2.2 Mediation analyses

Based on our SEM we tested whether inflammation, hypertension, diabetes, and serum HDL and triglycerides levels mediated the relationship between obesity and WMH volume. Sample size for these analyses after outlier exclusion was 11,803. This parallel mediation model was calculated in lavaan package for R (v. 0.6-5). Next, we investigated whether volume of subcortical structures mediated the relationship between obesity measures and cognition. We focused on subcortical structures as they were significantly related to WMH volume in the SEM. Finally, we tested whether WMH volume mediated the relationship between obesity measures and grey matter morphometry, but also obesity measures and cognition. In all mediation analyses that included cognitive variables we used education as a covariate of no interest.

In each analysis, criteria for a significant mediation were: 1) significant total effect; 2) significant mediation effect; 3) total and mediation effects of the same sign (Baron and Kenny, 1986). Unless otherwise specified, mediation estimates and p-values were assessed using 1,000 quasi-Bayesian Monte Carlo simulations using ‘mediation’ package for R (Tingley et al., 2014). P-values were corrected for multiple comparisons using FDR Benjamini-Hochberg correction and were considered significant at α=0.05 (Benjamini and Hochberg, 1995). The correction was applied separately for each grey matter morphometry modality and collectively for all cognitive measures.

##### 2.6.2.3 Follow-up analysis on white matter microstructure

As a follow-up analysis, we decided to investigate the relationship between WMH load and white matter microstructure measures (FA and MD) for each WM tract from the John Hopkins University atlas (Mori et al., 2005; Wakana et al., 2007) to pinpoint spatial locations where WMH influence white matter microstructure. To this end, we used general linear model analysis with each WM parcel’s FA and MD values as outcome measures and WMH load as a predictor variable. P-values were corrected for multiple comparisons using FDR Benjamini-Hochberg correction and were considered significant at α=0.05 (Benjamini and Hochberg, 1995). Correction was applied separately for FA and MD analyses. Sample size for these analyses after outlier exclusion was 12,192.

## 3 Results

### 3.1 Structural equation model

With this model we tested overall associations between obesity, inflammation, dyslipidemia, hypertension, diabetes, cerebrovascular disease, grey matter morphometry and cognition. In line with our hypotheses, we found that obesity was related to increased inflammation, as measured by serum CRP, but also hypertension, diabetes diagnosis, and dyslipidemia (serum triglycerides and serum HDL). Inflammation, hypertension, diabetes, and dyslipidemia in turn, led to increased cerebrovascular disease, which further significantly affected cortical volume and thickness (negatively) and subcortical volumes (positively). Finally, we found that obesity, lower subcortical volumes and higher cerebrovascular disease were all directly related to impaired cognitive function (Table 1, Figure 1). Overall, 22% of the variance in inflammation, 13% in hypertension, 6% in diabetes, 54% in dyslipidemia, 26% in cerebrovascular disease, 33% in subcortical volumes, 10% in cortical thickness, 11% in cortical volume, and 25% of variance in cognition were explained by our model. Loadings of latent variables for this model can be found in Table S4. Note that increased scores on the latent variable ‘cognition’ means worse cognitive performance. The SEM estimation produced the following fit measures: Χ^2^(11721)=257,582 p<0.0001; SRMR=0.066, RMSEA=0.058, CFI=0.912.

**Table 1.**
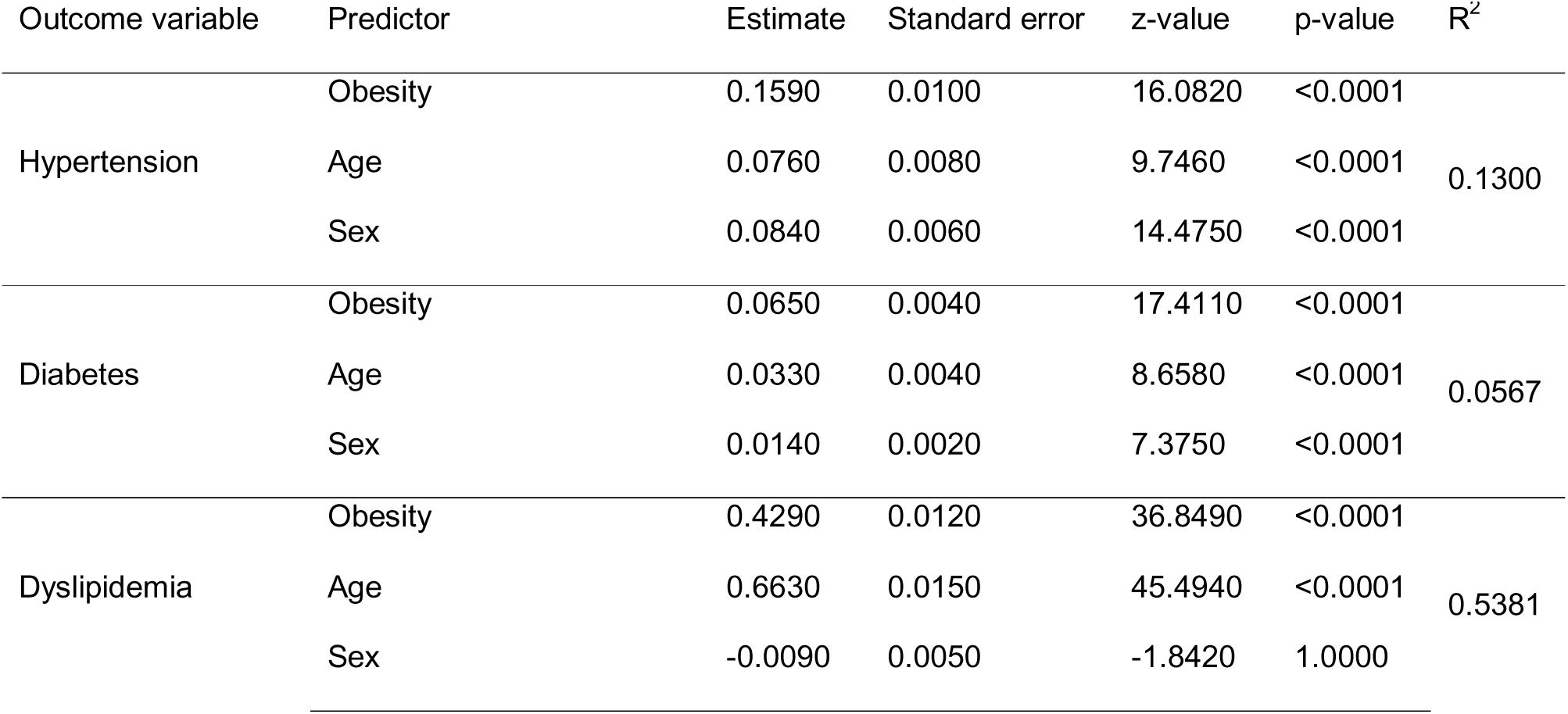
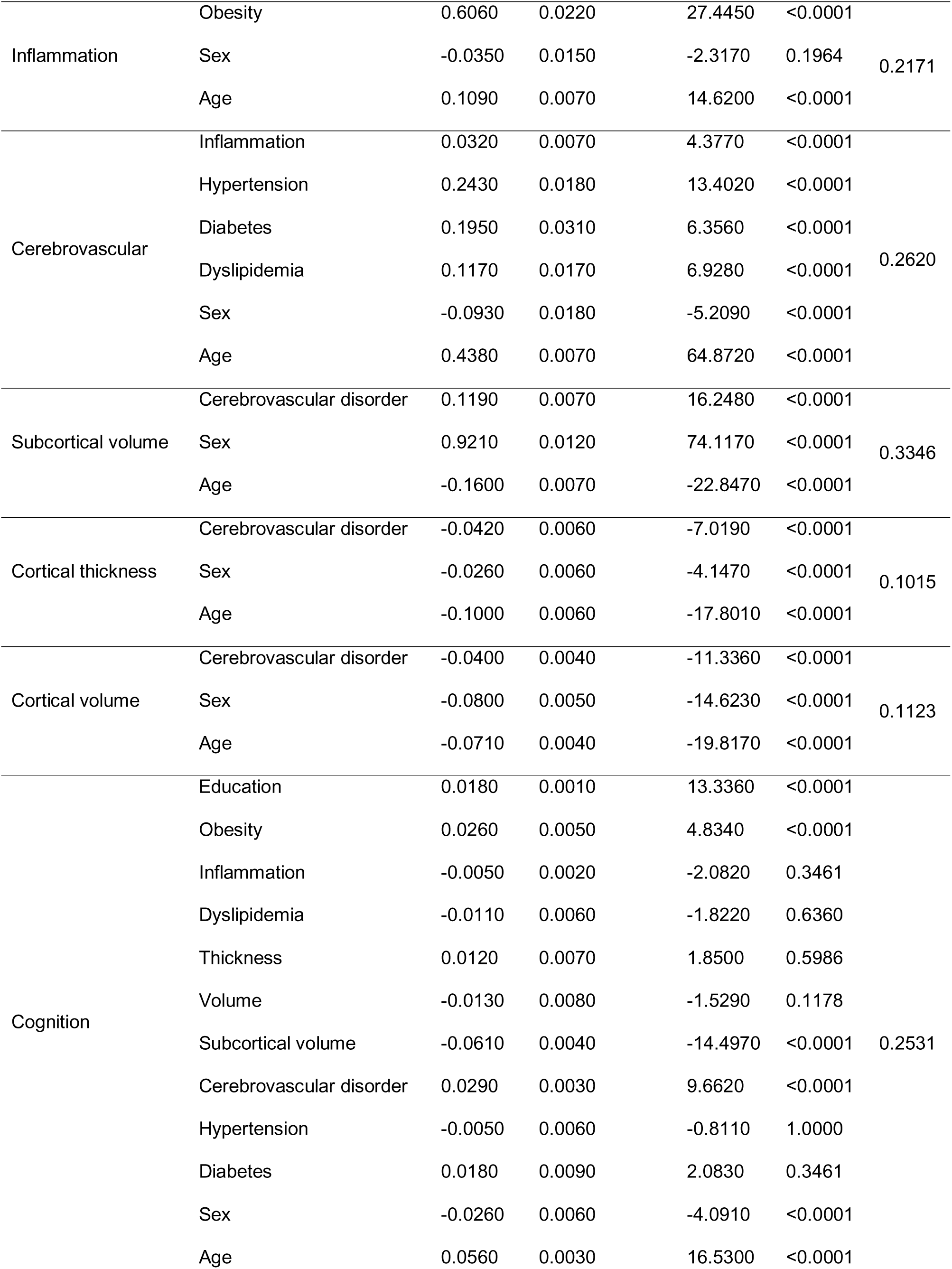
Statistics of the structural equation model

### 3.2 Relationship between obesity and cerebrovascular disease is mediated by inflammation, hypertension and diabetes

We found that the relationship between WHR and WMH load was mediated by levels of CRP, however, CRP levels did not mediate the relationship between BMI and WMH (Table 2). The relationships between both BMI and WHR, and WMH load were mediated by hypertension and diabetes (Table 2), but not by serum HDL or triglyceride levels. Overall, inflammation, hypertension, and diabetes mediated approximately 20% of the relationships between BMI and WMH, and WHR and WMH.

**Table 2.**
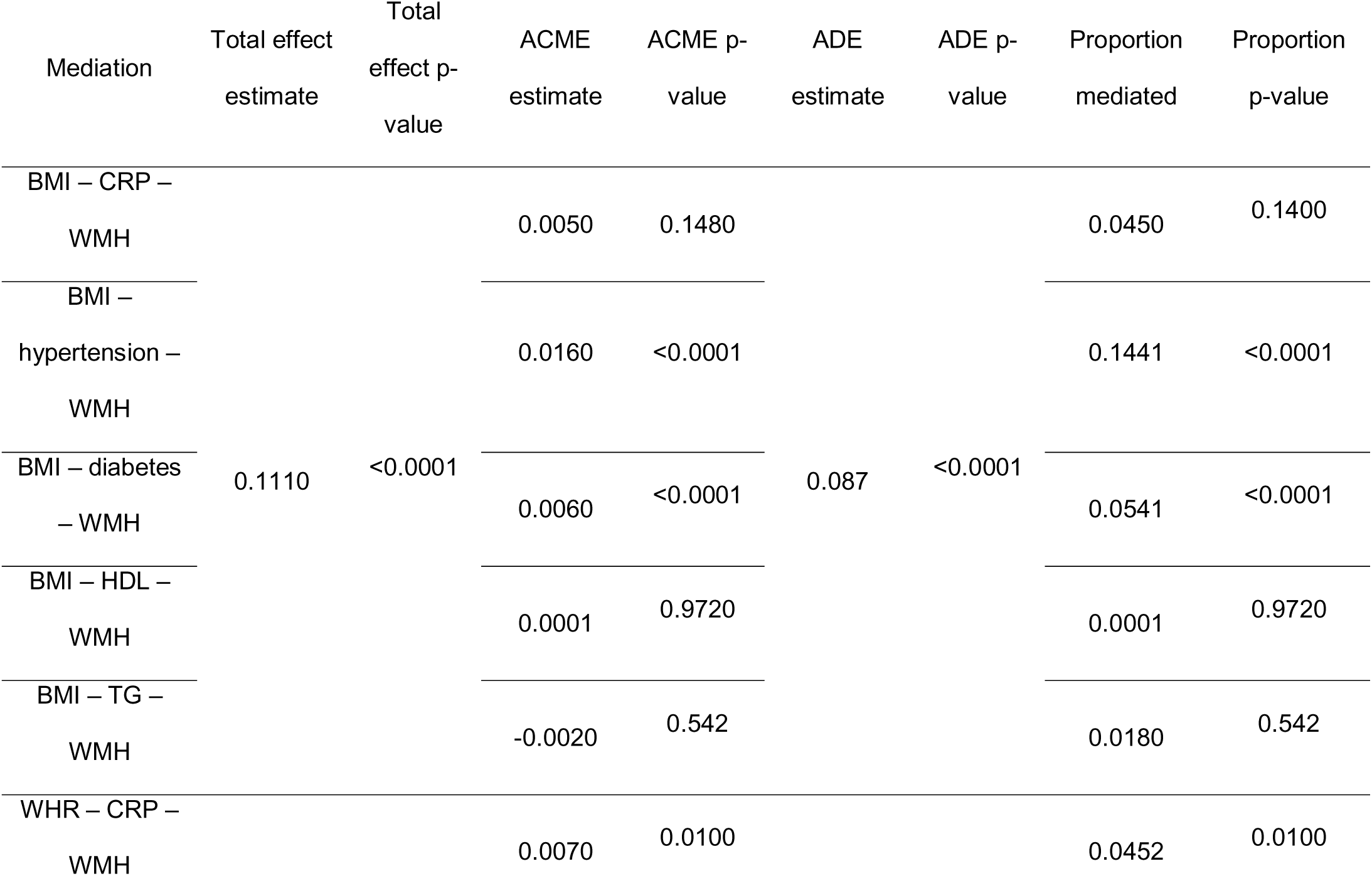
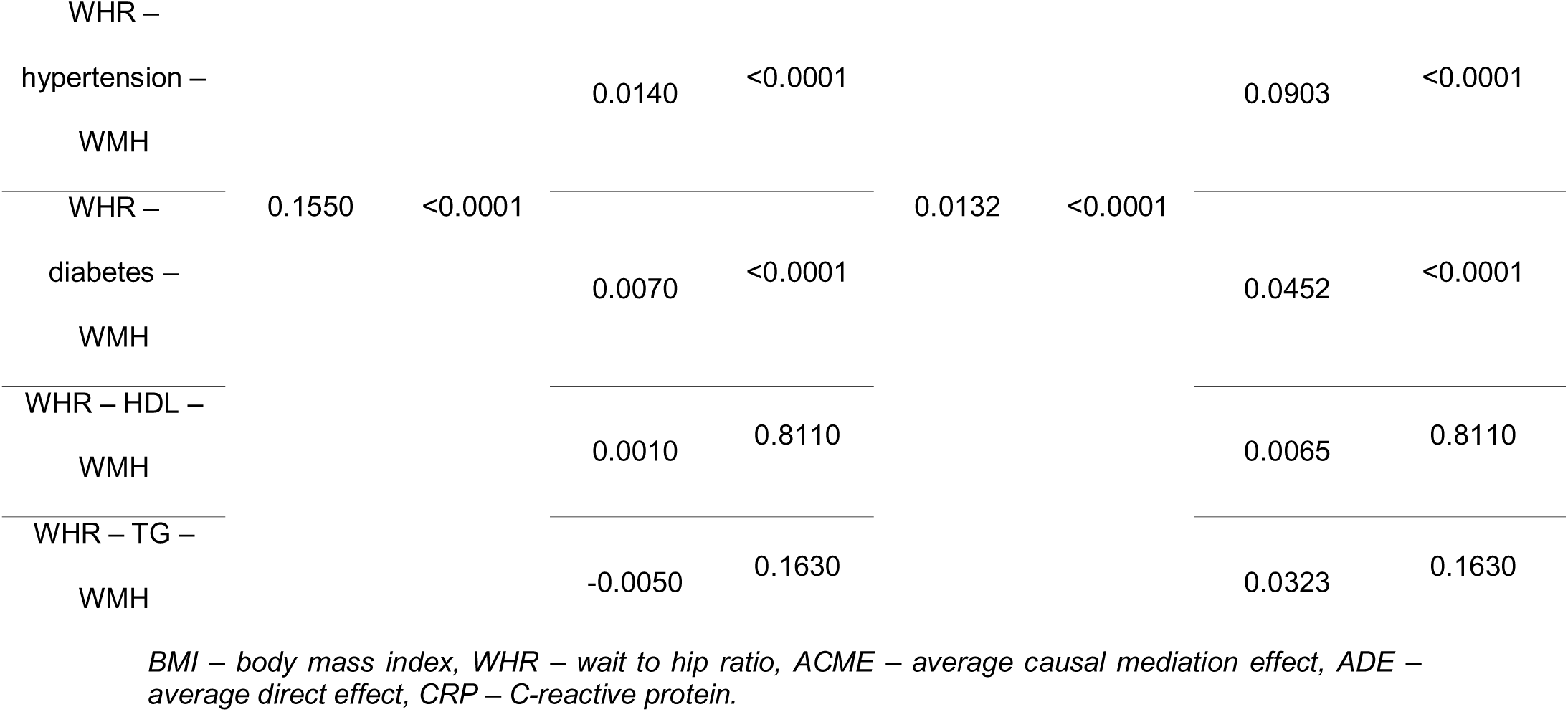
Mediation analysis – inflammation (blood C-reactive protein levels), hypertension and diabetes mediate the relationship between waist-to-hip ratio and white matter hyperintensities load.

### 3.3 Obesity is associated with grey matter alterations and cognitive impairment

We observed widespread associations between obesity measures, cortical thickness and grey matter volumes. Regarding cortical thickness, we found negative correlations with BMI and WHR in the bilateral temporal, entorhinal, orbitofrontal and cingulate cortices. Positive correlations were also observed with the frontal, parietal and occipital cortex, which were more pronounced for BMI than for WHR (Table S5, Figure 2). Similarly, BMI and WHR correlated negatively with grey matter volume in the bilateral temporal, temporoparietal and parietal cortices, with less pronounced positive correlations in the occipital and posterior cingulate cortex (Table S5, Figure 2). With regard to subcortical structures, BMI correlated negatively with volume of the left thalamus and bilateral caudate, putamen, pallidum, hippocampus, amygdala and nucleus accumbens (Table S6, Figure 2). WHR correlated negatively with volume of the left thalamus and hippocampus, and bilateral putamen, hippocampus, amygdala, and nucleus accumbens. WHR also correlated positively with the right thalamus volume (Table S6, Figure 2).

**Figure 2.**
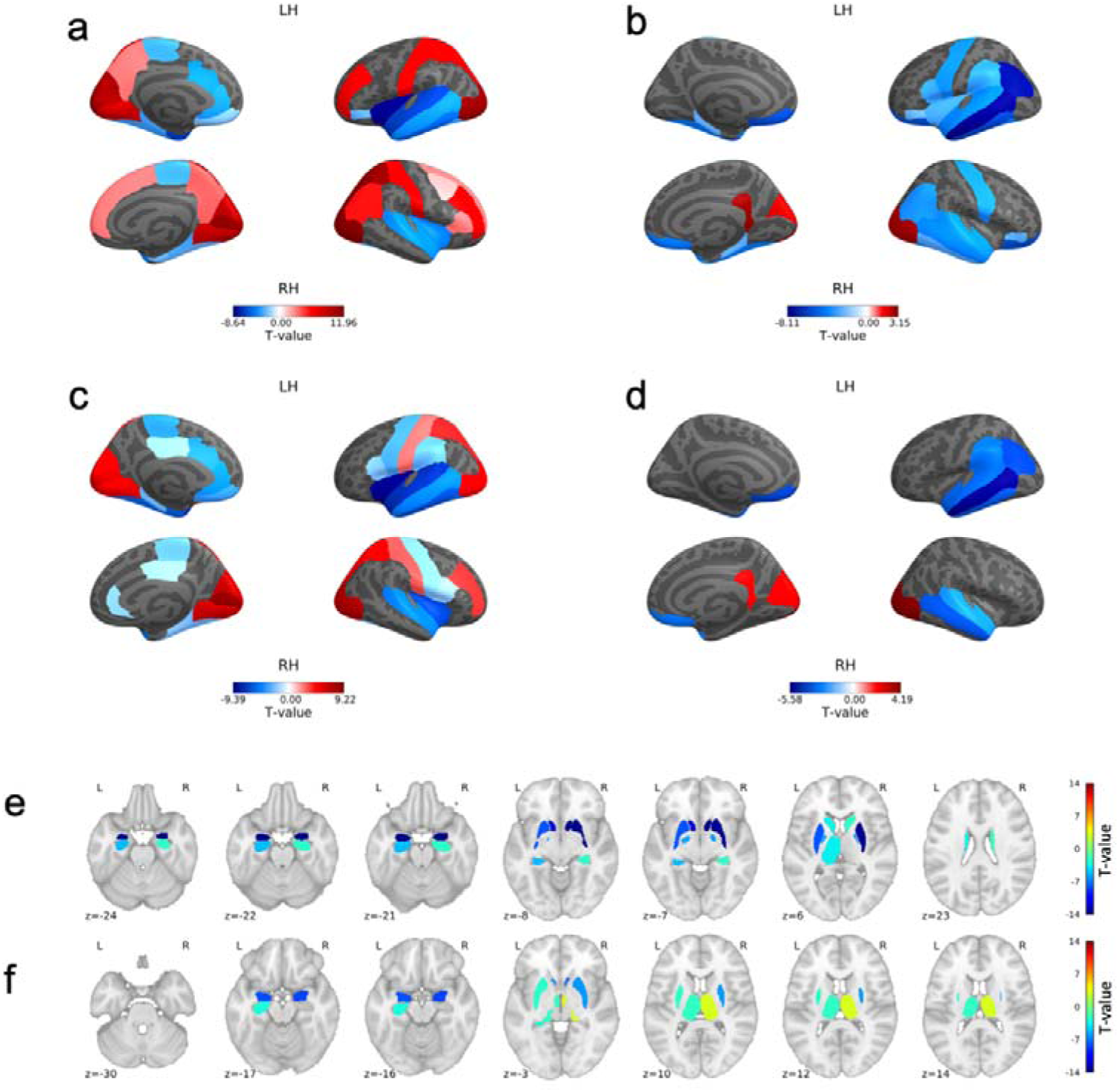
Associations between: **a** body mass index and cortical thickness; **b** body mass index and cortical volume; **c** waist-to-hip ratio and cortical thickness; **d** waist-to-hip ratio and cortical volume; **e** body mass index and subcortical volume**; f** waist-to-hip ratio and subcortical volume. Figures depict T-values of significant associations. RH – right hemisphere, LH – left hemisphere.

Regarding cognition, BMI was related to lower working memory and better executive functions and visuospatial memory (Table S7), while WHR was related to lower working memory, fluid intelligence, prospective memory, and higher reaction time (Table S7).

#### 3.3.1 Volume of subcortical structures mediates the relationship between obesity and cognition

None of the associations between BMI and cognition were mediated by volume of subcortical structures. However, consistent with our SEM, some associations between high WHR, and poorer cognition were mediated by volumes of subcortical structures (Table S8). Here, the bilateral amygdala mediated the associations between WHR and prospective memory (6%), and WHR and reaction time (left: 13%, right: 10%). Finally, the bilateral nucleus accumbens and left hippocampus were significant mediators between WHR and reaction time (accumbens left: 8%, right: 9%, hippocampus: 3%). Proportions of the relationships mediated were small, suggesting that factors other than volumes of subcortical structures likely influence cognition.

#### 3.3.2 Cerebrovascular disease mediates the relationship between obesity and cognition, and obesity and grey matter morphometry

The association between BMI and working memory was mediated in 19% by WMH load (Table 3). The relationships between WHR and working memory, fluid intelligence, prospective memory and reaction time were mediated by WMH load (Table 3). Proportions mediated ranged from 11% for working memory to 16% for fluid intelligence.

**Table 3.**
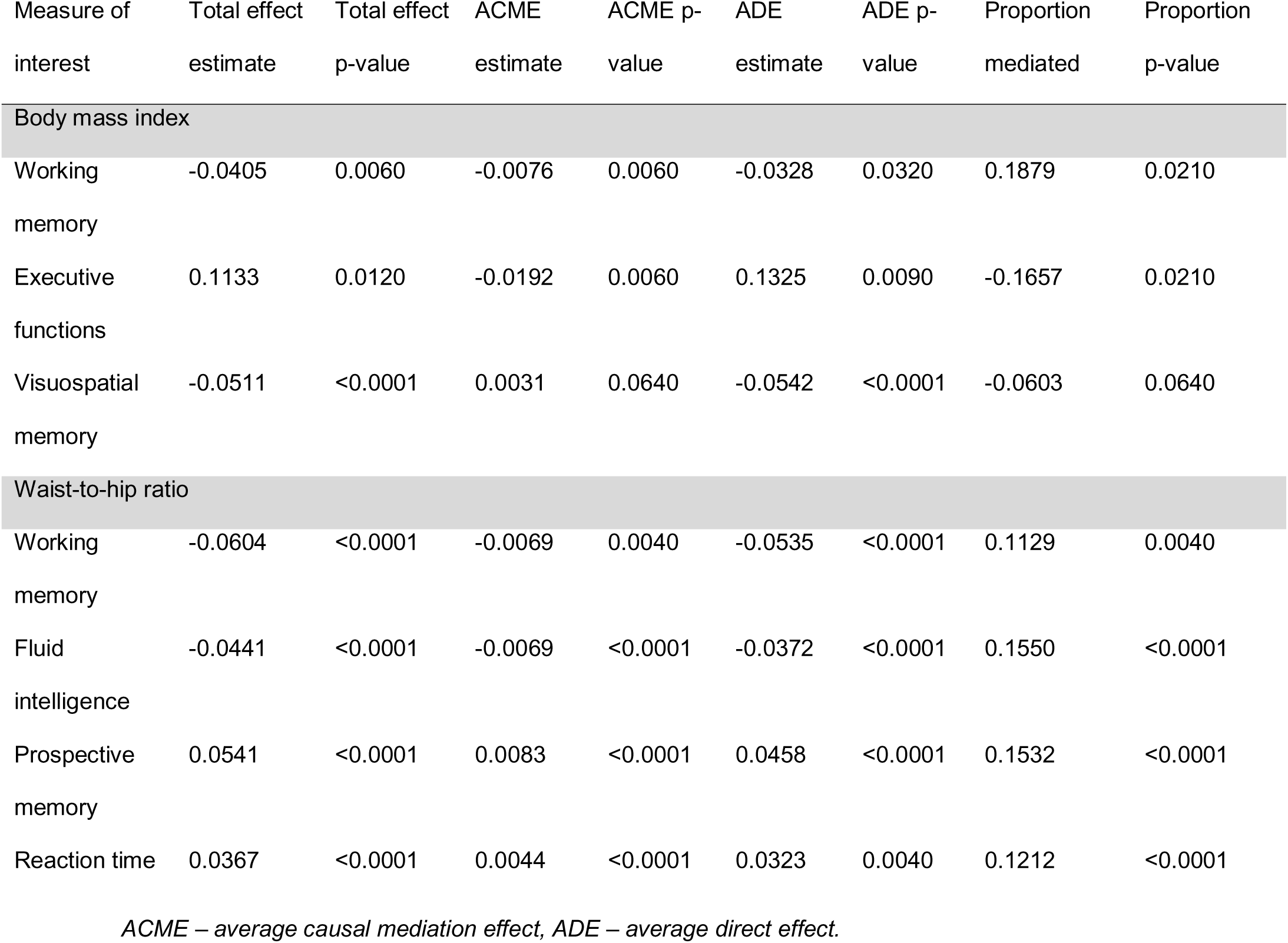
Mediation effects of white matter hyperintensities on the associations between body mass index and cognitive abilities.

| Measure of interest | Total effect estimate | Total effect p-value | ACME estimate | ACME p-value | ADE estimate | ADE p-value | Proportion mediated | Proportion p-value |
| --- | --- | --- | --- | --- | --- | --- | --- | --- |
| Body mass index |  |  |  |  |  |  |  |  |
| Working memory | -0.0405 | 0.0060 | -0.0076 | 0.0060 | -0.0328 | 0.0320 | 0.1879 | 0.0210 |
| Executive functions | 0.1133 | 0.0120 | -0.0192 | 0.0060 | 0.1325 | 0.0090 | -0.1657 | 0.0210 |
| Visuospatial memory | -0.0511 | <0.0001 | 0.0031 | 0.0640 | -0.0542 | <0.0001 | -0.0603 | 0.0640 |
| Waist-to-hip ratio |  |  |  |  |  |  |  |  |
| Working memory | -0.0604 | <0.0001 | -0.0069 | 0.0040 | -0.0535 | <0.0001 | 0.1129 | 0.0040 |
| Fluid intelligence | -0.0441 | <0.0001 | -0.0069 | <0.0001 | -0.0372 | <0.0001 | 0.1550 | <0.0001 |
| Prospective memory | 0.0541 | <0.0001 | 0.0083 | <0.0001 | 0.0458 | <0.0001 | 0.1532 | <0.0001 |
| Reaction time | 0.0367 | <0.0001 | 0.0044 | <0.0001 | 0.0323 | 0.0040 | 0.1212 | <0.0001 |

*ACME – average causal mediation effect, ADE – average direct effect.*

Further, WMH mediated the relationship between obesity measures (both BMI and WHR) and cortical thickness and volume in all DKT parcels that were negatively related to obesity measures (Figure 3, Table S9, Table S10, Table S11, Table S12). These parcels were predominantly located in the bilateral temporal lobes, entorhinal cortex, temporoparietal junction, cingulate cortex, and orbitofrontal cortex. The magnitude of mediation varied between 9% and 68%, with the largest mediating effect in the paracentral lobule and insula. With regard to subcortical structures, WMH mediated the relationship between BMI and subcortical volumes in the bilateral amygdala and the left nucleus accumbens (Figure 3, Table S13). WMH also mediated the relationship between WHR and volume of the bilateral amygdala and nucleus accumbens, and the right thalamus (Figure 3, Table S13). The mediations were modest ranging from 4%-15%.

**Figure 3.**
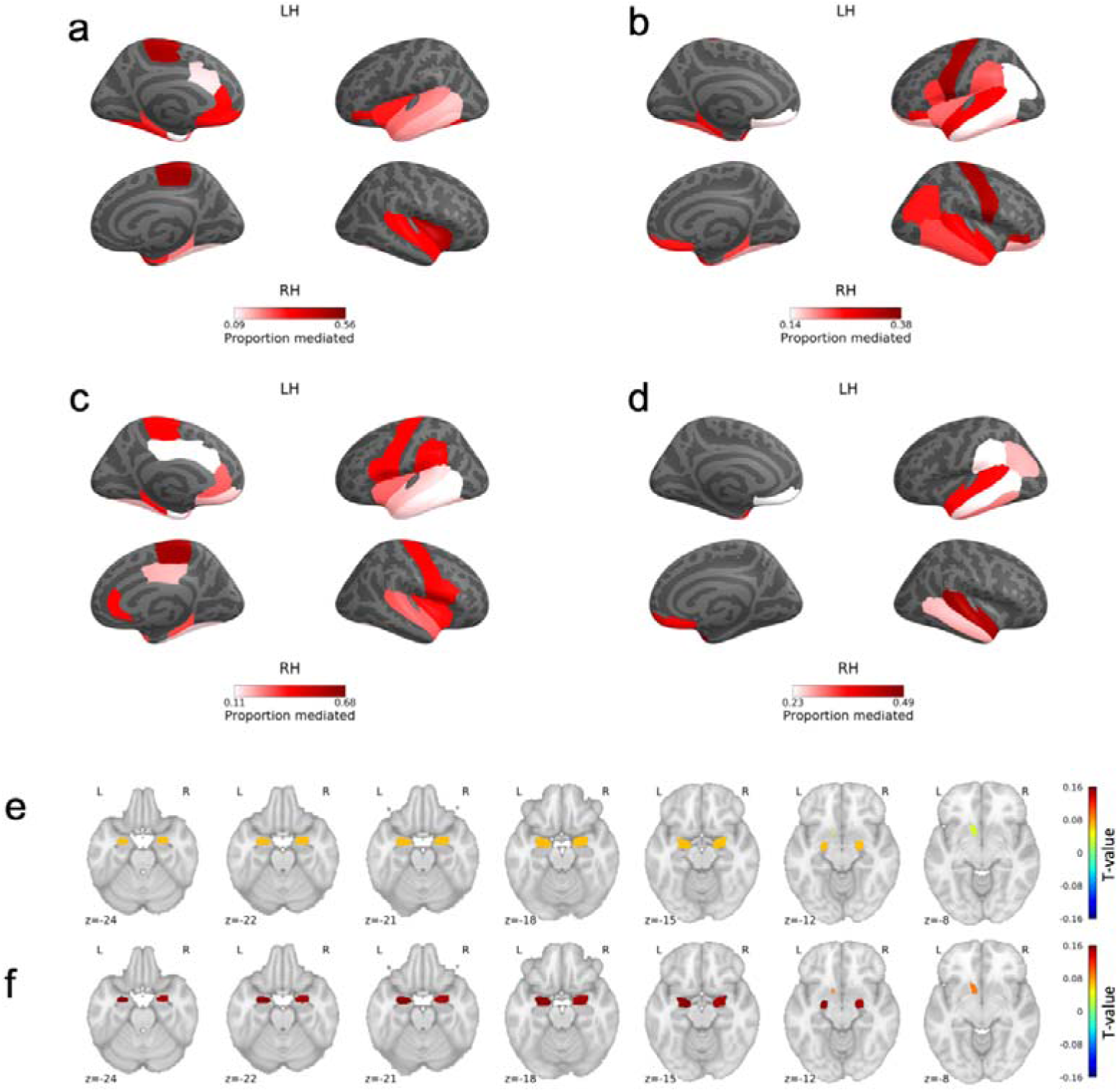
Proportion of relationships mediated by white matter hyperintensities between **a** body mass index and cortical thickness; **b** body mass index and cortical volume; **c** waist-to-hip ratio and cortical thickness; **d** waist-to-hip ratio and cortical volume; **e** body mass index and subcortical volumes**; f** waist-to-hip ratio and subcortical volumes. Figures depict only significant consistent mediations. RH – right hemisphere, LH – left hemisphere.

### 3.4 Relationship between white matter hyperintensities and white matter microstructure

We found significant associations between WMH load and FA. WMH were related to lower FA, most notably in the posterior thalamic radiation and anterior corona radiata (Figure 4, Table S14). WMH were also associated with predominantly higher mean diffusivity with the strongest associations in the entire corona radiata, posterior thalamic radiation and the superior frontooccipital fasciculus (Figure 4, Table S14). Relationships between MD and WMH were organised in a superior-to-inferior gradient, where superior white matter tracts showed more positive associations, while inferior tracts showed more negative associations.

**Figure 4.**
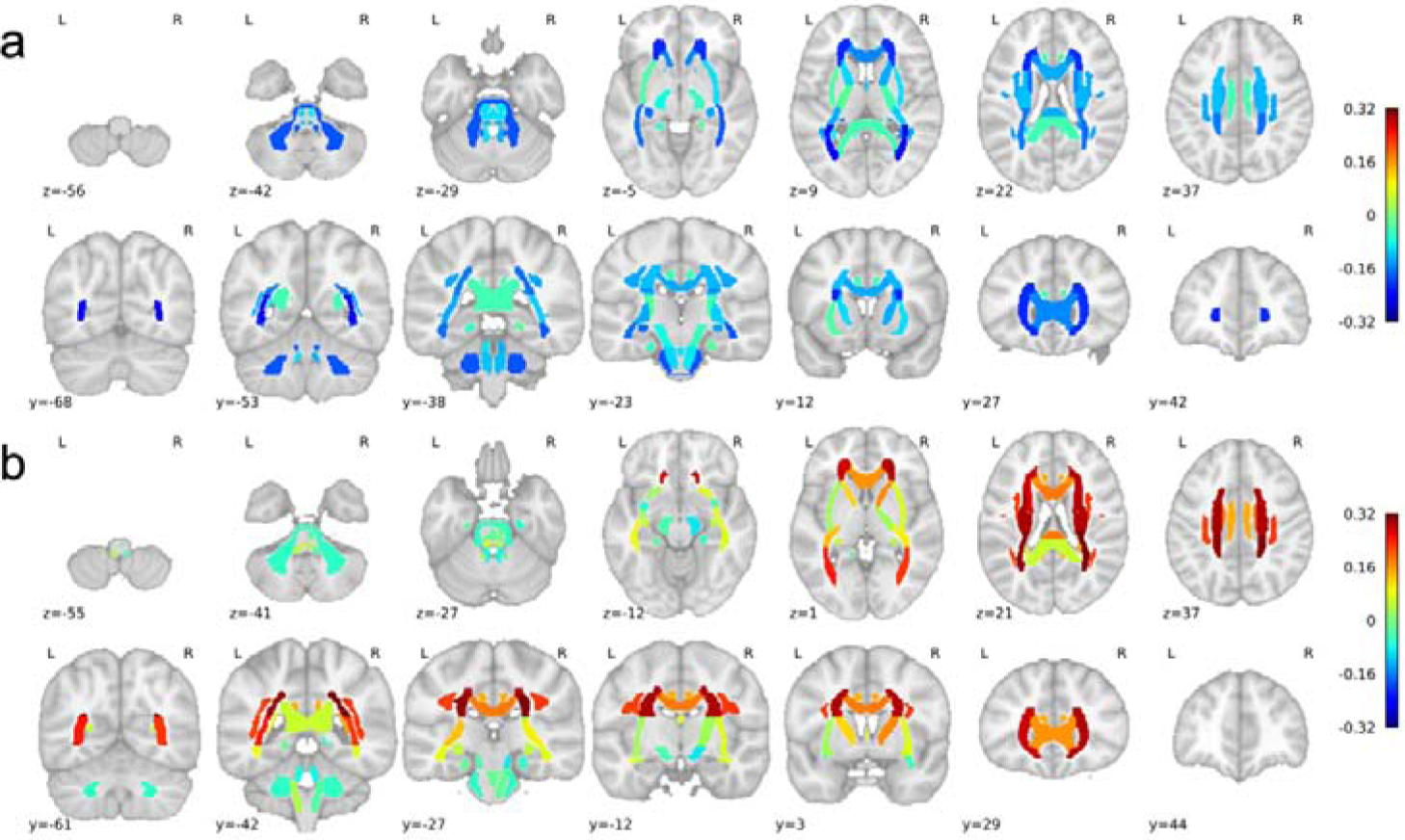
Significant associations between white matter hyperintensities and: **a** fractional anisotropy; **b** mean diffusivity. R – right, L – left.

## 4 Discussion

We investigated the relationship between obesity, grey and white matter disruption, and cognition, and the possible mechanisms that link them. Using structural equation modelling, we find that obesity is related to systemic inflammation, hypertension, diabetes and dyslipidemia. In turn, inflammation, hypertension and diabetes lead to cerebrovascular disease, which is further related to grey matter alterations and impaired cognition. We describe specific spatial patterns of cortical atrophy that are associated with obesity and white matter hyperintensities. We also find that certain cognitive domains are related to obesity, white matter hyperintensities and subcortical volumes. Finally, we show that white matter hyperintensities are related to lower fractional anisotropy and higher mean diffusivity in the majority of white matter tracts. Coupled with the fact that white matter hyperintensities affect cognition independently of cortical atrophy, this implies a role of structural connectivity disruptions in poor cognition in obesity.

The associations between obesity, especially visceral adiposity, and systemic inflammation, dyslipidemia, hypertension, and diabetes, are consistent with previous reports (Mathieu et al., 2009). The same factors lead to white matter lesions and changes in FA and MD within white matter tracts (Castellon and Bogdanova, 2016; Van Dijk et al., 2005; Kuo et al., 2010; Swardfager et al., 2017; Tamura and Araki, 2015; Zhao et al., 2019). Our mediation analyses are in line with those findings and show that inflammation, hypertension, and diabetes mediate the relationship between obesity and WMH independent of each other. Consequently, they might separately constitute potential targets for therapeutic interventions aimed at improving brain and cognitive outcomes in obesity.

Further, we showed that WMH are related to cortical grey matter loss. In our SEM, cerebrovascular disease was related to lower cortical thickness and volume. Our follow-up analyses show that WMH mediate the relationship between obesity and grey matter volume and thickness loss, but also that obesity is negatively related to grey matter morphometry measures independent of WMH. This suggests that the metabolic abnormalities due to adiposity may damage grey matter both directly and via white matter lesions. The spatial pattern of reductions in cortical volume is in line with previous reports. In their meta-analysis, Garcia-Garcia and colleagues showed that higher BMI was related to lower cortical volume in the medial prefrontal cortex, precentral gyrus, temporal pole and cerebellum (García-García et al., 2018), which resembles our findings. With regard to cortical thickness, our results are also consistent with previous reports showing lower thickness in the temporal and occipital lobes in obesity (Beyer et al., 2019b; Medic et al., 2016; Shaw et al., 2018; Vainik et al., 2018; Veit et al., 2014). Contrary to our hypotheses, in the SEM, we showed that the volume of WMH was related to higher volume of subcortical structures. This may represent true hypertrophy of these structures, but it may also be explained by subcortical segmentation inaccuracies, where periventricular WMH (which appear hypointense on T1w MRI, with similar intensities to the neighboring subcortical grey matter structures) are classified as grey matter, leading to inflated estimated of subcortical volumes. This phenomenon, in fact, was demonstrated in a recent study (Dadar et al., 2020). Irrespective of this, our analyses of the direct relationship between obesity and subcortical volumes produced results consistent with the previous literature (Beyer et al., 2019b; Dekkers et al., 2019; García-García et al., 2019; Hamer and Batty, 2019), which shows that that obesity is related to lower volume of subcortical structures.

While some studies report associations between obesity and global cortical thickness/grey matter volume (Dekkers et al., 2019; Gustafson et al., 2004; Hamer and Batty, 2019; Ward et al., 2005), our analysis allowed a more fine-grained spatial differentiation. Thus, we also identified areas with greater cortical thickness and volume associated with higher BMI predominantly in the occipital and frontal lobes. Similar findings were present in some previous studies regarding cortical volumes in young adults and using smaller sample sizes (Herrmann et al., 2019; Pannacciulli et al., 2006; Taki et al., 2008), but we are aware of only one study that showed similar patterns in cortical thickness, in this case using the Human Connectome Project dataset (Vainik et al., 2018). One explanation for higher cortical thickness and volume in obesity might be neuroinflammation-related astrocytosis and microgliosis which could result in higher volume and thickness of grey matter regions (DiSabato et al., 2016; Guillemot-Legris and Muccioli, 2017; Heneka et al., 2015; Hsuchou et al., 2012b; Maldonado-Ruiz et al., 2017; Streit et al., 1999). Alternatively, it is possible that juxtacortical white matter damage in obesity might be incorrectly classified as grey matter and therefore grey matter would appear thicker (Chard et al., 2002, 2010; Dadar et al., 2020). Further, given the possible bidirectional relationship between brain and obesity (García-García et al., 2019), it is also possible that higher cortical thickness and volume in obesity in certain areas reflects a brain-based trait that leads to over-eating and obesity, while lower volumes in other areas may reflect an effect of obesity and metabolic syndrome, as suggested by our SEM. Higher cortical thickness in frontal and parietal regions is in line with findings showing obesity-related functional and anatomical alterations in the same regions that are ultimately linked to eating behaviour (Han et al., 2018; Kakoschke et al., 2019; Mehl et al., 2019; Wang et al., 2017). In keeping with the theory that certain cortical alterations represent an underlying phenotype related to over-eating, Vainik et al. showed that cortical thickness associations with BMI in young adults were largely heritable (Vainik et al., 2018). Nonetheless, the main direction of associations here are consistent with tissue damage, as demonstrated by the mediating effect of metabolic syndrome and cerebrovascular lesions.

Our findings relating obesity to grey matter atrophy in the temporoparietal regions support previous theories that obesity accelerates aging and is a risk factor for dementia (Alford et al., 2018; Ronan et al., 2016). Indeed, aging is associated with loss of cortical grey matter in the temporal lobes and temporoparietal regions first, while other areas are affected at a later time (Fjell et al., 2009). Additionally, brain atrophy in Alzheimer’s disease also seems to follow this pattern (Singh et al., 2006). Specifically, differences in cortical thickness between individuals with mild cognitive impairments and healthy individuals were found in the temporal, parietal and frontal cortices, while individuals with Alzheimer’s disease showed further cortical thinning in the lateral temporal lobe. This resembles the obesity-related atrophy patterns found here in the temporal, parietal and frontal cortices, and provides a mechanism for the known association between adiposity in midlife and later Alzheimer’s disease (Kivipelto et al., 2005; Whitmer et al., 2007).

Finally, we show that obesity affects cognition via white matter hyperintensities, volume of subcortical structures and other independent mechanisms. Consistent with the idea that it is visceral adipose tissue that is most related to metabolic syndrome (Mathieu et al., 2009), we find that WHR, rather than BMI, is a better correlate of poor cognition, as it was associated with worse outcomes in multiple cognitive domains while BMI was only related to lower working memory. According to our follow-up analysis, one mechanism in which obesity, and hence WMH, affects cognition is via white matter dysconnectivity. We show that WMH were associated with white matter microstructural changes (as measured by diffusion weighted imaging) in tracts related to cognition, such as the corona radiata or thalamic radiations (Bendlin et al., 2010; Benito-León et al., 2017; Hua et al., 2014; Zhang et al., 2018). In this respect, our research replicates the study by Zhang and colleagues showing that white matter microstructure mediates the relationship between obesity and cognition (Zhang et al., 2018).

Our study is cross-sectional. A longitudinal design would allow for better causal inferences on the origin of brain changes and impaired cognitive performance in obesity. Further, only CRP blood levels were used to quantify inflammation. Extending the methodology to include more inflammatory factors is warranted and recommended for future studies on the topic.

In sum, based on our results, a model of obesity-related brain and cognitive changes emerges. Greater adiposity causes low-grade systemic inflammation, hypertension and diabetes, which in turn lead to small vessel disease occurring as white matter hyperintensities. Together, they cause subcortical and cortical alterations and cognitive deficits in obesity, which can be partially ascribed to white matter dysconnectivity. Our results suggest that obesity can lead to accelerated brain aging and act as a risk factor for dementia. This has clinical implications for the management of obesity and prevention of dementia. In fact, studies show that lifestyle interventions including dietary changes and anti-hypertensive medication might be effective in decreasing the pace of cognitive deficits in older adults (Debette and Markus, 2010; Dufouil et al., 2005; Lam et al., 2015; Lee et al., 2014; de Leeuw et al., 2002; Ngandu et al., 2015).

## 5 Authors contributions

All authors contributed to each stage of the presented study

## 8 Supplementary Tables

**Table S1.**
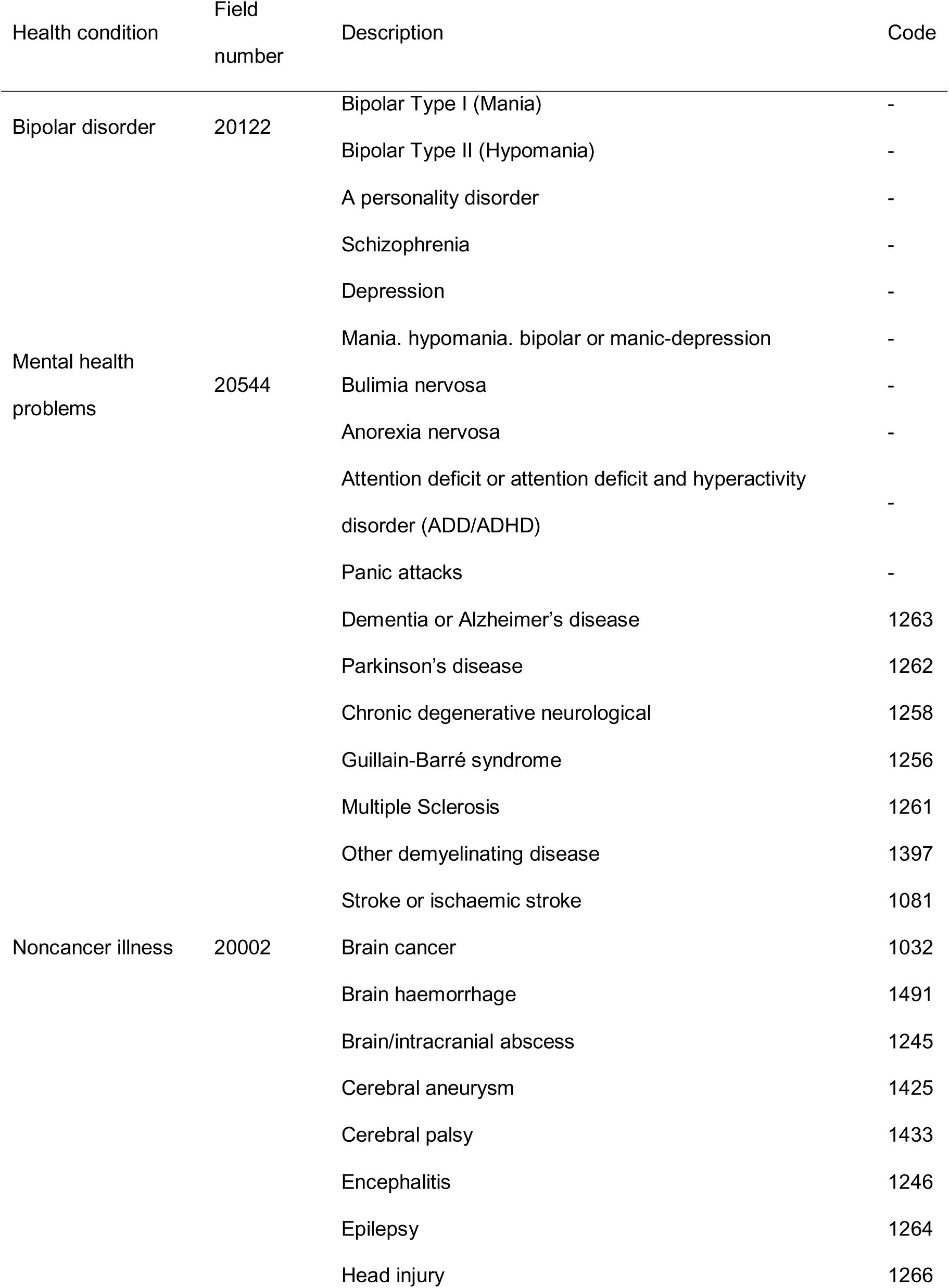
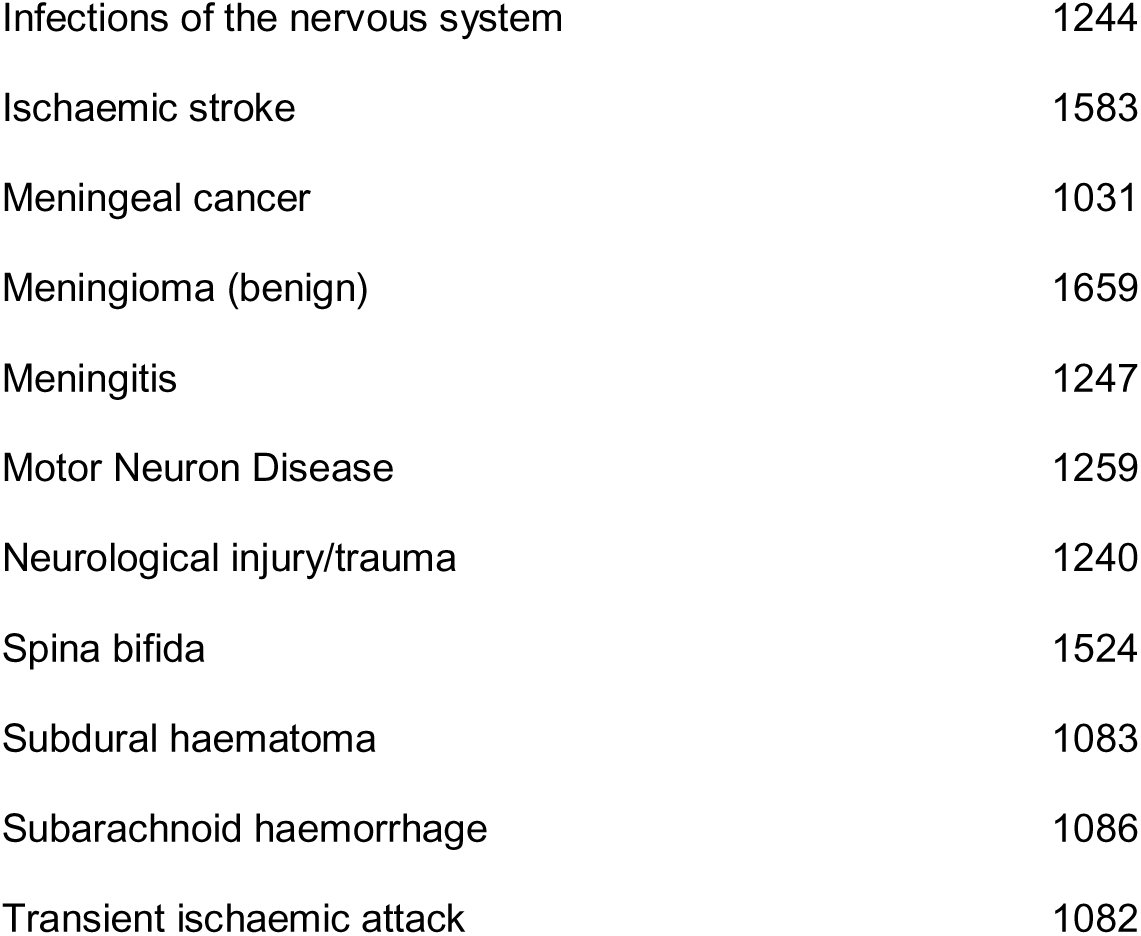
UK Biobank field IDs used for exclusion of participants

**Table S2.** UK Biobank field IDs used in this study.

| Field name | Field number | Description |
| --- | --- | --- |
| Body mass index | 21001 |  |
| Waist circumference | 48 |  |
| Hip circumference | 49 |  |
| Age | 21003 |  |
| Sex | 31 |  |
| Qualifications/education | 6138 |  |
| Total volume of white matter hyperintensities | 25781 |  |
| C-reactive protein blood level | 30710 |  |
| Triglycerides blood level | 30870 |  |
| High density lipoprotein level | 30760 |  |
| Cortical thickness | 27174-27204; 27267-27297 |  |
| Cortical volume | 27205-27235; 27298-27328 |  |
| Diabetes diagnosed by doctor | 2443 |  |
| Vascular / heart problems diagnosed by doctor | 6150 | High blood<br>pressure |
| Subcortical volumes | 25011 - 25024 |  |
| Fractional anisotropy | 25056 - 25103 |  |
| Mean diffusivity | 25104 - 25151 |  |
| Working memory – maximum digits remembered<br>correctly | 4282 |  |
| Fluid intelligence score | 20016 |  |
| Executive functions – numbers of puzzles correct | 21004 |  |
| Prospective memory | 20018 |  |
| Visuospatial memory – number of incorrect matches | 399 |  |
| Reaction time | 404 |  |

**Table S3.** Sample characteristics

| Characteristic | SEM sample |  | Post-SEM sample |  |
| --- | --- | --- | --- | --- |
|  | Mean | Standard deviation | Mean | Standard deviation |
| Age [years] | 63 | 8 | 63 | 7 |
| Body mass index [kg/m<sup>2</sup>] | 26.32 | 3.87 | 26.44 | 4.09 |
| Waist-to-hip ratio | 0.87 | 0.084 | 0.87 | 0.084 |
|  | Distribution |  | Distribution |  |
| Sex | 8191 ♀ 8247 ♂ |  | 7609 ♀ 7653 ♂ |  |
| Hypertension | 4047 (24.62%) |  | 3708 (24.3%) |  |
| Diabetes | 832 (5.06%) |  | 743 (4.9%) |  |

**Table S4.**
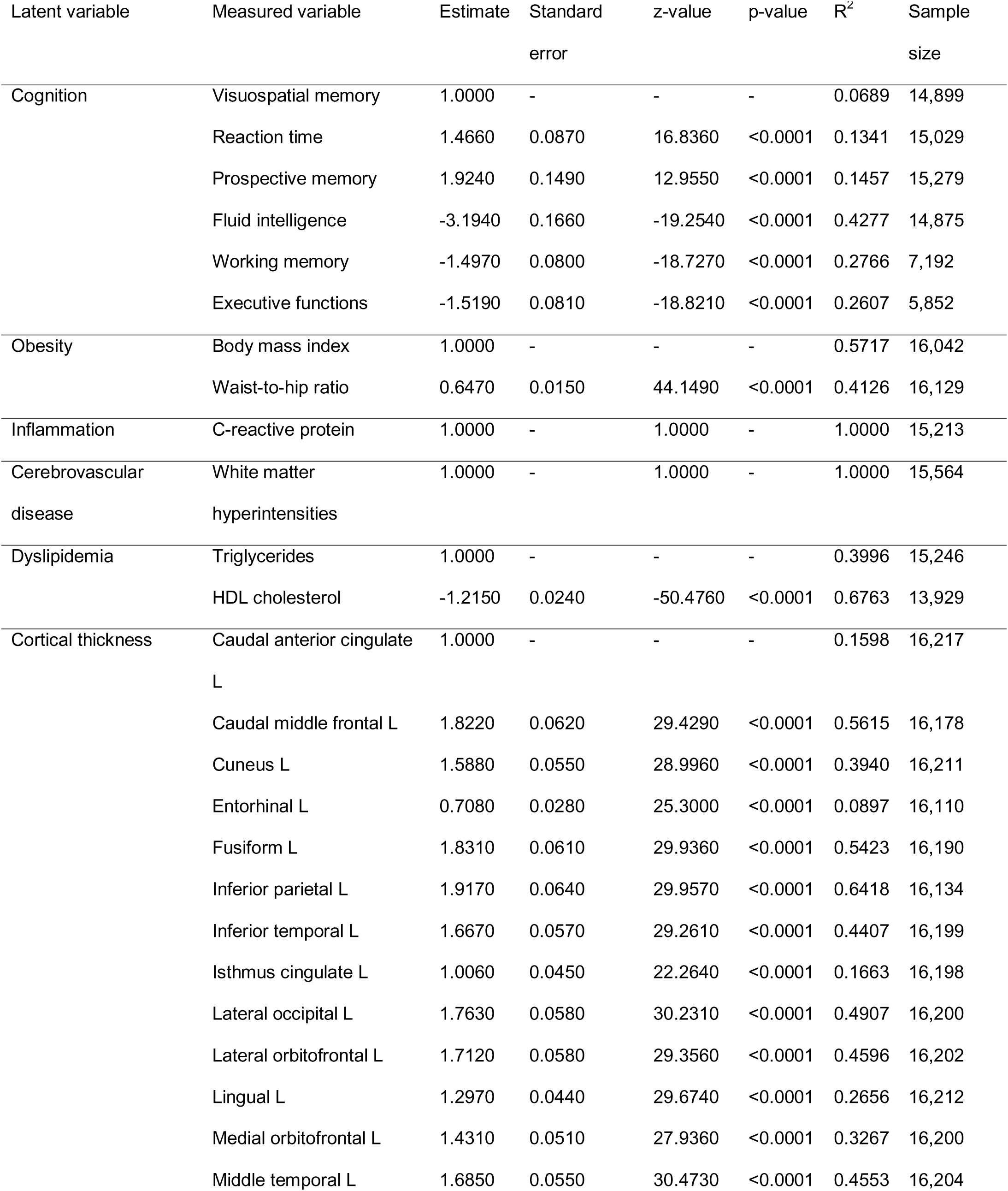
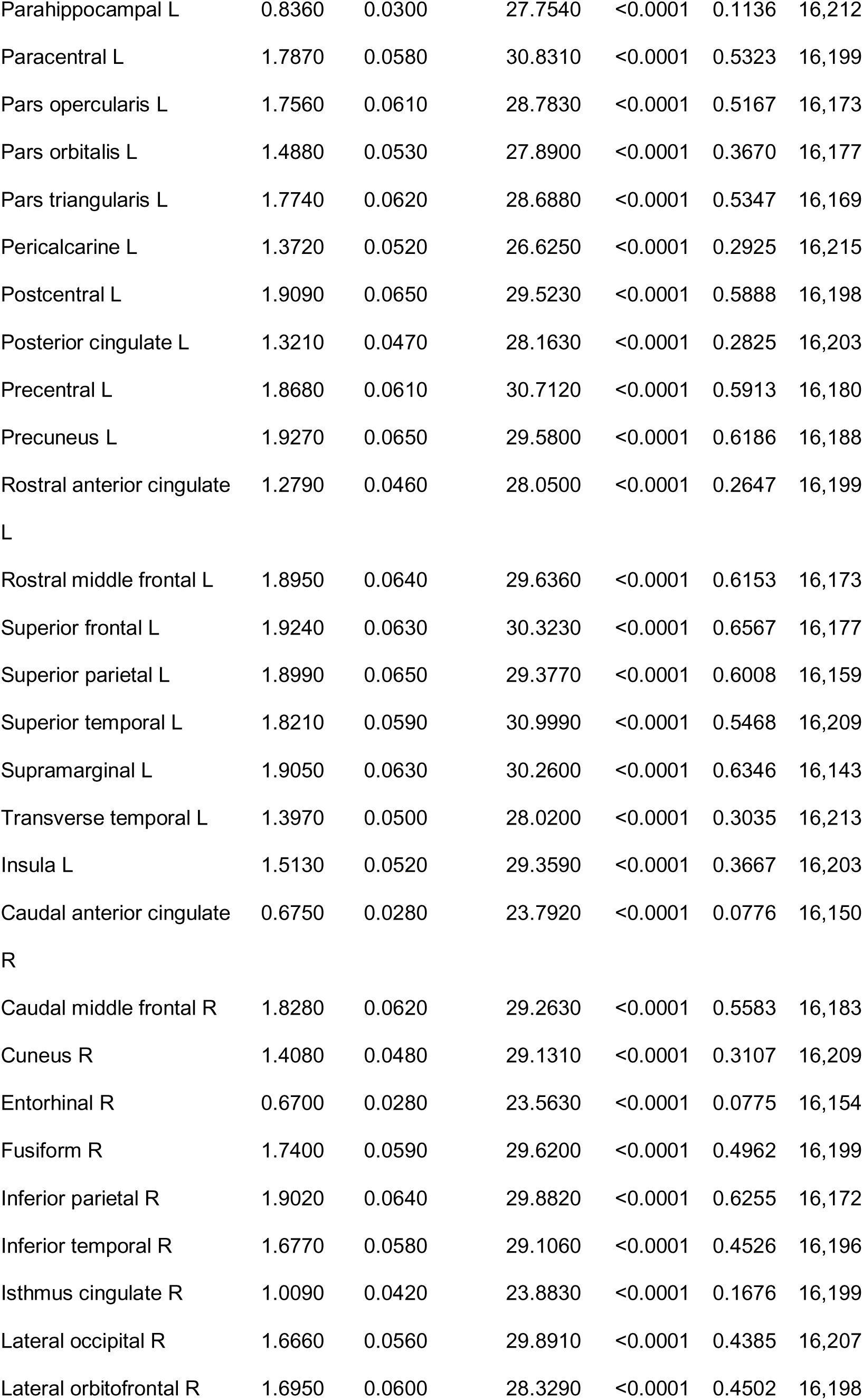
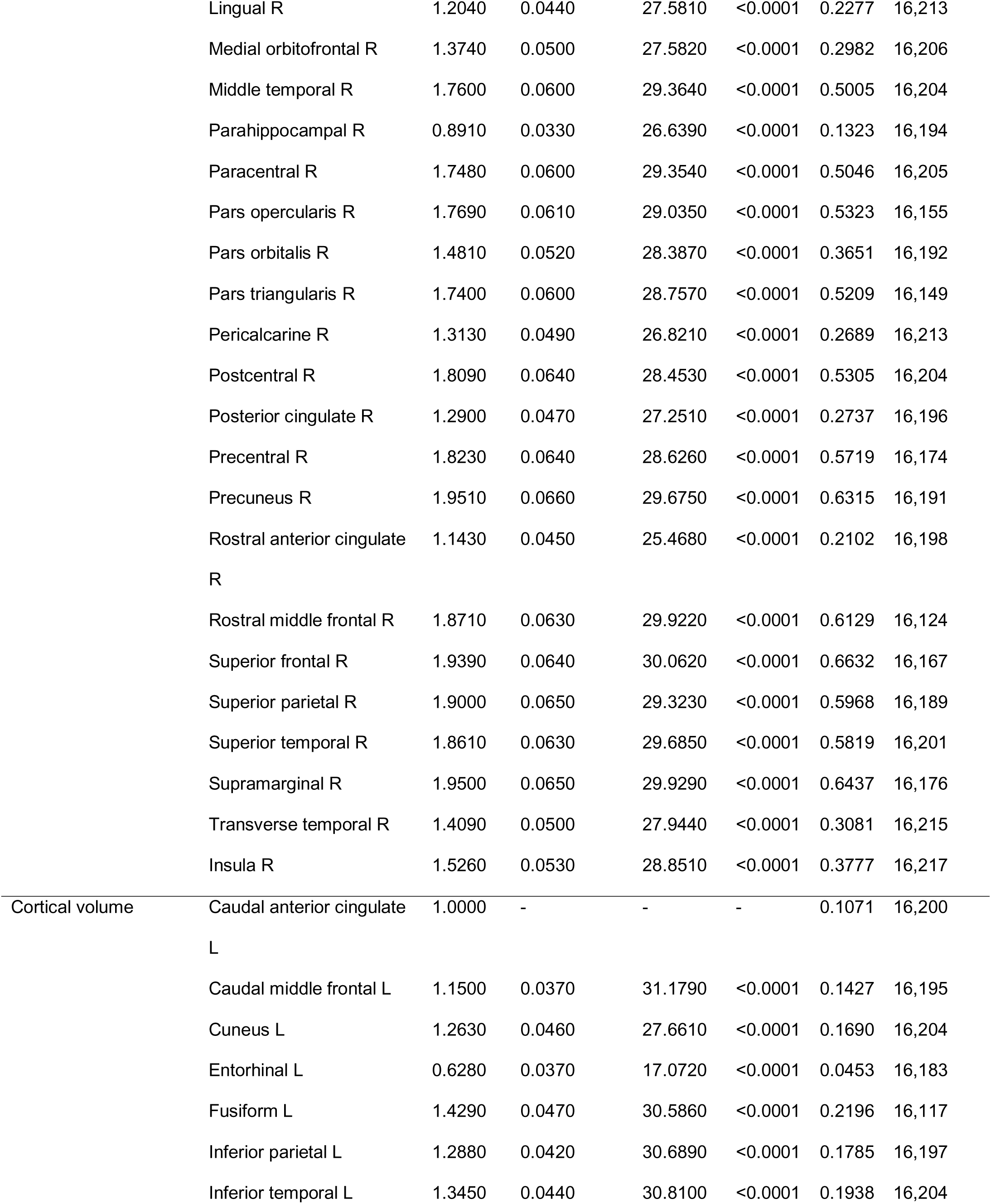
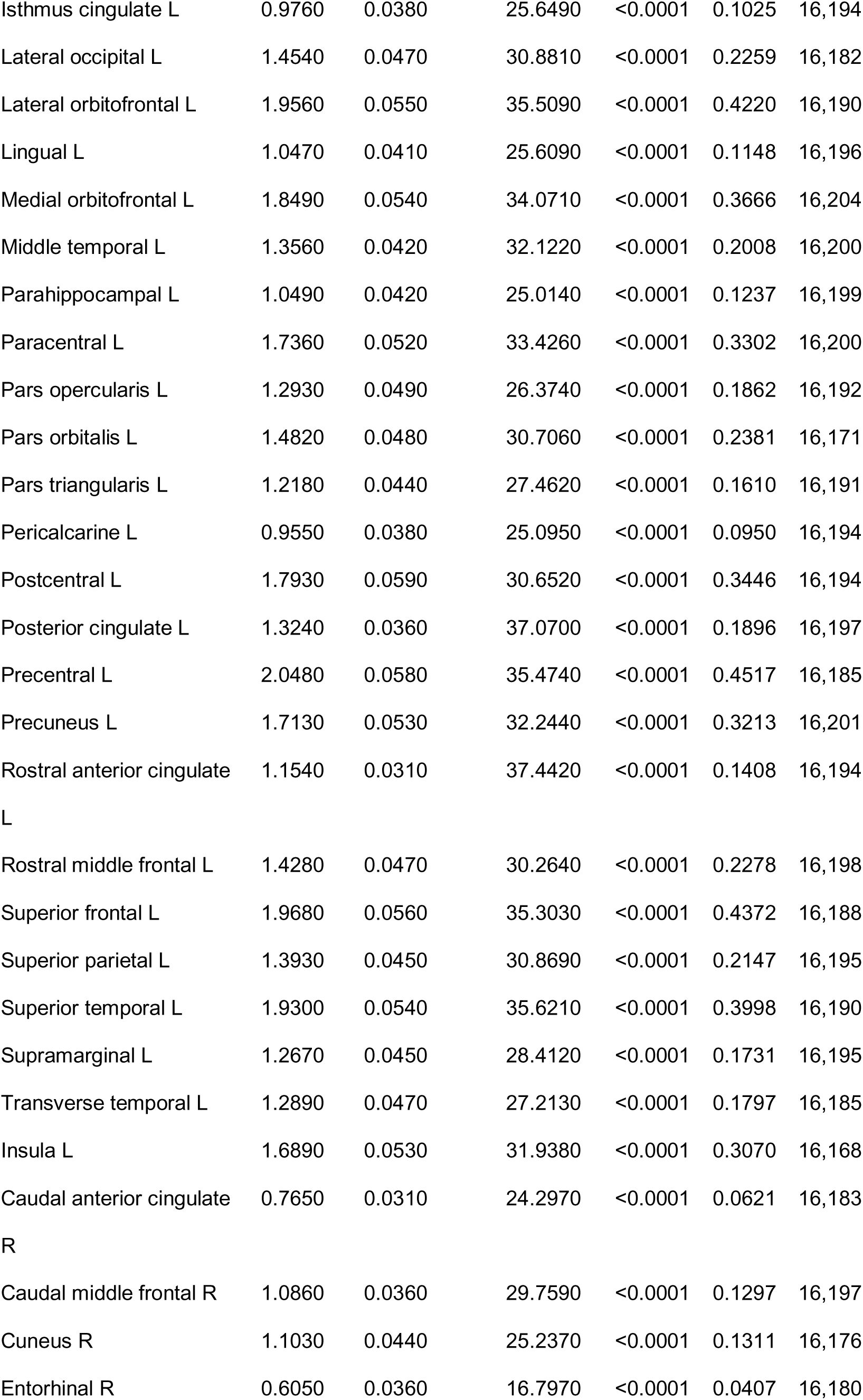
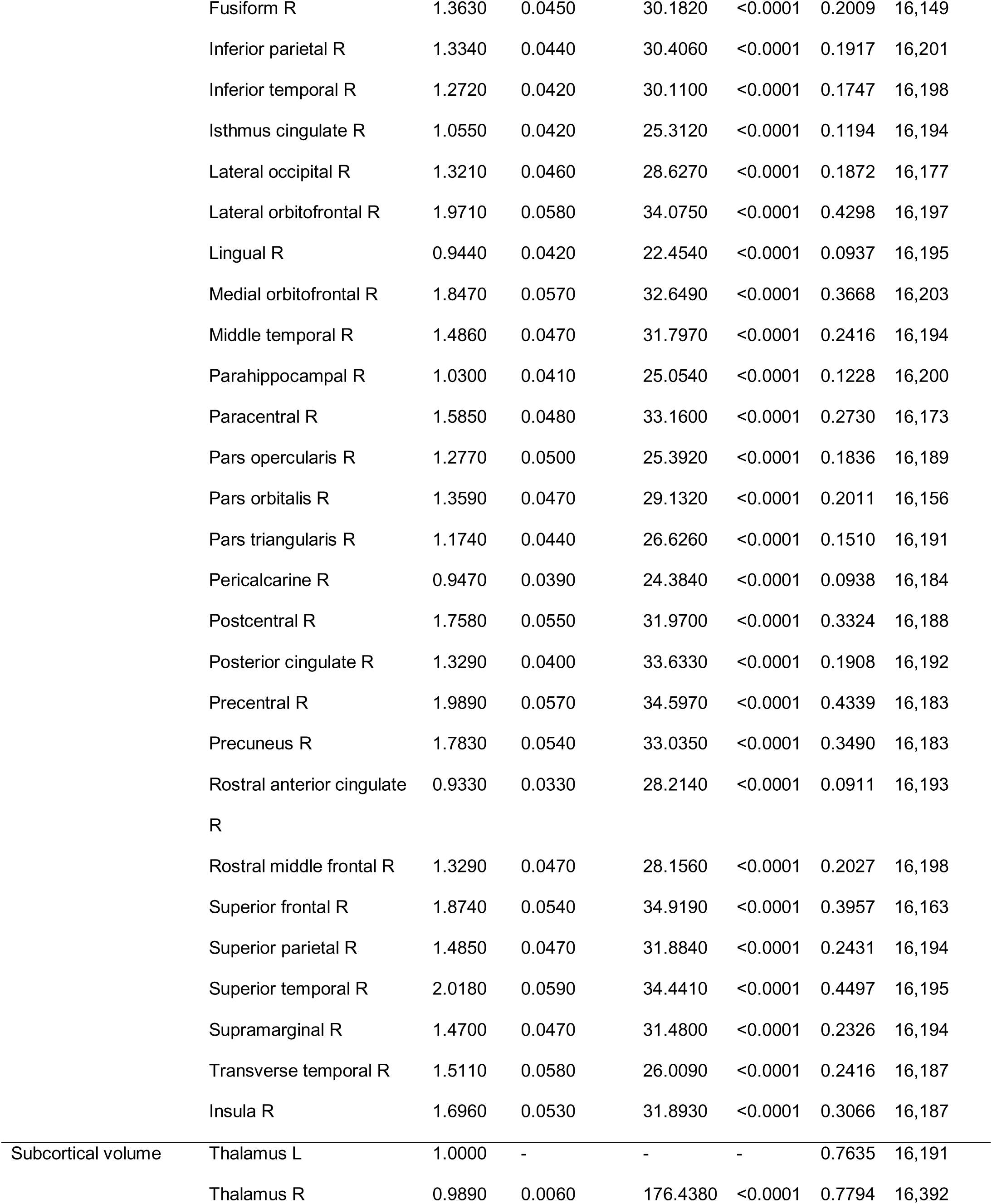
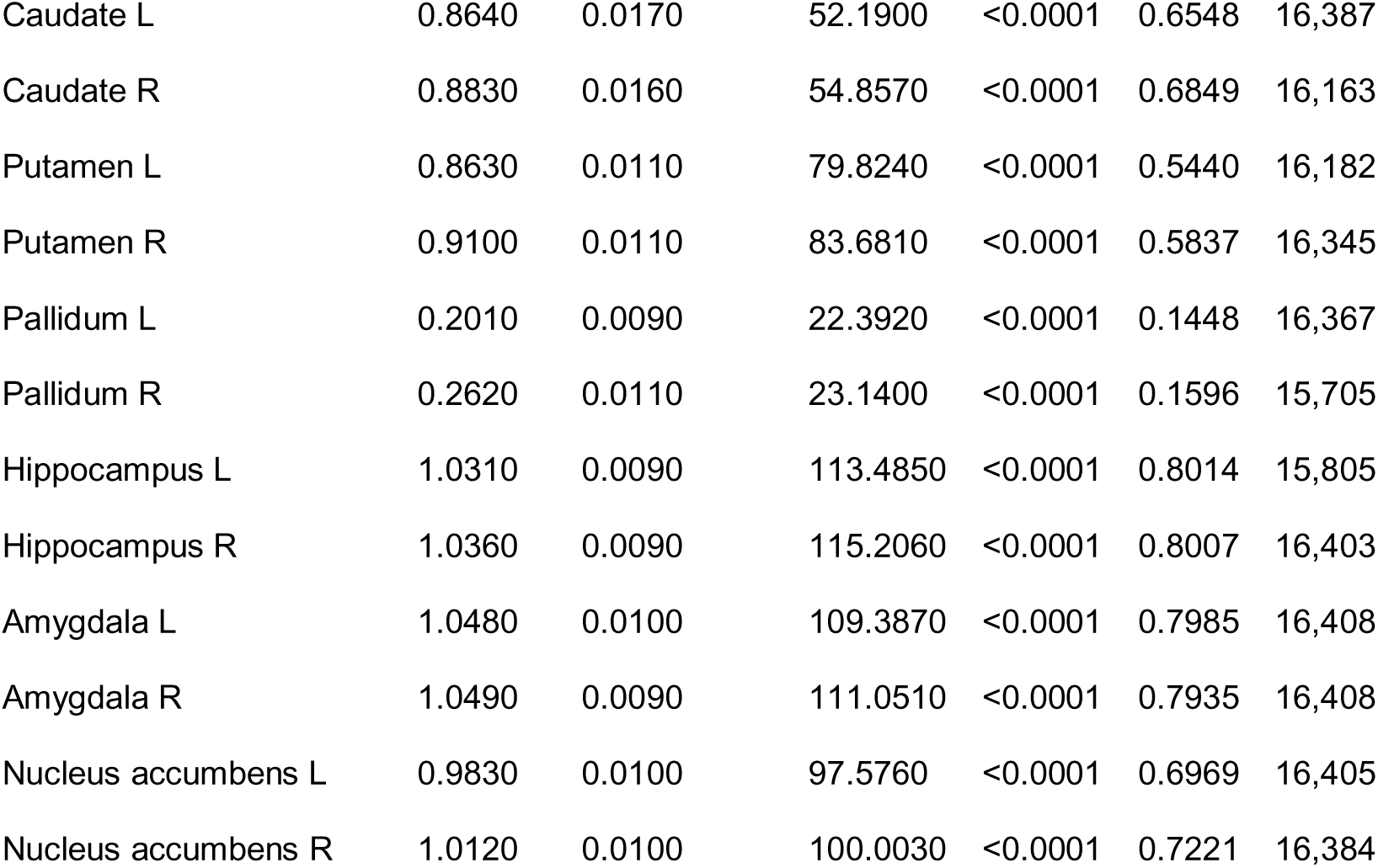
Loadings, standard deviations and p-values of measured variables on latent variables.

**Table S5.** Associations between body mass index and waist-to-hip ratio for each DKT parcel for cortical thickness and volume measures.

| Parcel | BMI |  |  |  | WHR |  |  |  |
| --- | --- | --- | --- | --- | --- | --- | --- | --- |
|  | Cortical thickness |  | Cortical volume |  | Cortical thickness |  | Cortical volume |  |
|  | p-value | T-value | p-value | T-value | p-value | T-value | p-value | T-value |
| Caudal anterior cingulate L | <0.0001 | -4.3258 | 0.9429 | 0.0907 | 0.0001 | -4.3086 | 0.6707 | -0.6613 |
| Caudal middle frontal L | 0.1981 | 1.4504 | 0.4481 | -1.0289 | 0.1949 | -1.4476 | 0.1955 | -1.8589 |
| Cuneus L | <0.0001 | 10.9913 | 0.3978 | 1.1192 | <0.0001 | 5.9231 | 0.7013 | 0.5914 |
| Entorhinal L | <0.0001 | -6.7839 | 0.7048 | -0.5677 | <0.0001 | -6.1648 | 0.7309 | -0.5058 |
| Fusiform L | 0.0020 | -3.2798 | 0.0002 | -4.2356 | <0.0001 | -5.6736 | 0.1079 | -2.1998 |
| Inferior parietal L | 0.0807 | 1.9219 | <0.0001 | -7.6794 | 0.6371 | 0.5271 | 0.0004 | -4.2760 |
| Inferior temporal L | <0.0001 | -5.3125 | <0.0001 | -5.9987 | <0.0001 | -6.5717 | 0.0012 | -3.8578 |
| Isthmus cingulate L | 0.6819 | -0.4707 | 0.1815 | 1.6472 | 0.2661 | -1.2528 | 0.4835 | 1.1144 |
| Lateral occipital L | <0.0001 | 11.8931 | 0.3543 | 1.2342 | <0.0001 | 6.3532 | 0.3424 | 1.4831 |
| Lateral orbitofrontal L | 0.2998 | -1.1828 | <0.0001 | -5.5587 | 0.0888 | -1.8798 | 0.4245 | -1.2858 |
| Lingual L | <0.0001 | 8.6943 | 0.1502 | 1.7560 | <0.0001 | 5.3852 | 0.5347 | 1.0143 |
| Medial orbitofrontal L | 0.0397 | -2.2520 | <0.0001 | -5.8404 | 0.0004 | -3.8093 | 0.0004 | -4.2356 |
| Middle temporal L | 0.0001 | -4.2302 | <0.0001 | -8.1136 | <0.0001 | -5.1246 | <0.0001 | -5.5761 |
| Parahippocampal L | 0.0013 | -3.4283 | 0.0289 | -2.5375 | 0.0130 | -2.7544 | 0.3642 | -1.4300 |
| Paracentral L | 0.0011 | -3.4852 | 0.1254 | -1.8908 | 0.0001 | -4.0646 | 0.3271 | -1.5277 |
| Pars opercularis L | 0.4354 | -0.8935 | 0.0039 | -3.2728 | 0.0209 | -2.5505 | 0.7365 | -0.4762 |
| Pars orbitalis L | 0.0117 | -2.7272 | 0.0424 | -2.3850 | 0.0860 | -1.9154 | 0.8219 | -0.2766 |
| Pars triangularis L | 0.0510 | 2.1438 | 0.1502 | -1.7512 | 0.6066 | -0.5957 | 0.7247 | -0.5469 |
| Pericalcarine L | <0.0001 | 10.7098 | 0.7048 | 0.5694 | <0.0001 | 6.9566 | 0.5669 | 0.9395 |
| Postcentral L | <0.0001 | 6.5604 | 0.2126 | -1.5405 | 0.0138 | 2.7246 | 0.5769 | 0.9100 |
| Posterior cingulate L | 0.2746 | -1.2465 | 0.9782 | 0.0273 | 0.0463 | -2.2248 | 0.8543 | 0.2189 |
| Precentral L | 0.3221 | -1.1149 | 0.0005 | -3.8836 | 0.0013 | -3.4840 | 0.1216 | -2.1283 |
| Precuneus L | 0.0011 | 3.5026 | 0.9365 | -0.1369 | 0.5722 | 0.6771 | 0.9439 | 0.0895 |
| Rostral anterior cingulate L | 0.0001 | -4.0846 | 0.5480 | -0.8457 | <0.0001 | -4.6055 | 0.1440 | -2.0357 |
| Rostral middle frontal L | <0.0001 | 7.1726 | 0.9365 | -0.1521 | 0.1873 | 1.4797 | 0.8219 | -0.2849 |
| Superior frontal L | 0.0612 | 2.0555 | 0.8110 | 0.3768 | 0.1241 | -1.7058 | 0.6484 | -0.7252 |
| Superior parietal L | <0.0001 | 8.6311 | 0.1345 | -1.8447 | 0.0002 | 4.0183 | 0.6677 | -0.6817 |
| Superior temporal L | <0.0001 | -6.6343 | 0.0002 | -4.1579 | <0.0001 | -7.7182 | 0.0026 | -3.6258 |
| Supramarginal L | 0.3197 | -1.1316 | 0.0022 | -3.4449 | 0.0056 | -3.0441 | 0.0055 | -3.3886 |
| Transverse temporal L | 0.5886 | 0.6547 | 0.7048 | 0.5977 | 0.5817 | -0.6496 | 0.4737 | 1.1638 |
| Insula L | <0.0001 | -8.6445 | 0.0073 | -3.0556 | <0.0001 | -9.3928 | 0.4140 | -1.3399 |
| Caudal anterior cingulate R | 0.8940 | 0.1332 | 0.4915 | 0.9525 | 0.1437 | -1.6244 | 0.6006 | -0.8328 |
| Caudal middle frontal R | 0.0302 | 2.3660 | 0.7048 | -0.6120 | 0.8987 | -0.1273 | 0.6998 | -0.6102 |
| Cuneus R | <0.0001 | 10.6454 | 0.0431 | 2.3644 | <0.0001 | 7.9264 | 0.0371 | 2.7177 |
| Entorhinal R | 0.0152 | -2.6292 | 0.7206 | -0.5127 | 0.0770 | -1.9738 | 0.5816 | 0.8868 |
| Fusiform R | 0.0169 | -2.5823 | 0.0002 | -4.1116 | 0.0200 | -2.5881 | 0.4166 | -1.3162 |
| Inferior parietal R | <0.0001 | 5.5525 | 0.0007 | -3.7461 | 0.2693 | 1.2341 | 0.2548 | -1.6722 |
| Inferior temporal R | 0.5948 | -0.6316 | 0.0158 | -2.7857 | 0.3525 | -1.0459 | 0.5347 | -1.0169 |
| Isthmus cingulate R | 0.6785 | -0.4906 | 0.0057 | 3.1474 | 0.0953 | -1.8381 | 0.0490 | 2.5665 |
| Lateral occipital R | <0.0001 | 11.7354 | 0.0158 | 2.7893 | <0.0001 | 7.7040 | 0.0004 | 4.1874 |
| Lateral orbitofrontal R | 0.1520 | 1.6080 | <0.0001 | -5.7160 | 0.4246 | 0.9098 | 0.1624 | -1.9621 |
| Lingual R | <0.0001 | 9.1977 | 0.1917 | 1.6063 | <0.0001 | 6.6915 | 0.2276 | 1.7670 |
| Medial orbitofrontal R | 0.2014 | -1.4301 | <0.0001 | -4.5495 | 0.0749 | -1.9964 | 0.0073 | -3.2759 |
| Middle temporal R | 0.6408 | 0.5552 | 0.0001 | -4.4098 | 0.0883 | -1.8928 | 0.0008 | -4.0097 |
| Parahippocampal R | 0.0024 | -3.2269 | 0.0217 | -2.6499 | 0.0303 | -2.3955 | 0.5669 | -0.9415 |
| Paracentral R | 0.0013 | -3.4282 | 0.3641 | -1.1879 | 0.0060 | -3.0124 | 0.6484 | 0.7215 |
| Pars opercularis R | 0.4354 | 0.8923 | 0.2507 | -1.4410 | 0.0463 | -2.2027 | 0.7309 | -0.5182 |
| Pars orbitalis R | 0.7733 | 0.3044 | 0.0203 | -2.6875 | 0.6066 | 0.5862 | 0.7365 | 0.4656 |
| Pars triangularis R | 0.0013 | 3.4073 | 0.7048 | -0.5538 | 0.6371 | 0.5155 | 0.4835 | 1.1262 |
| Pericalcarine R | <0.0001 | 11.9597 | 0.1502 | 1.7557 | <0.0001 | 9.2168 | 0.0353 | 2.7650 |
| Postcentral R | <0.0001 | 7.9700 | 0.9429 | 0.1014 | 0.0042 | 3.1368 | 0.7748 | 0.3855 |
| Posterior cingulate R | 0.1970 | -1.4649 | 0.0829 | -2.0813 | 0.0463 | -2.2136 | 0.0746 | -2.3651 |
| Precentral R | 0.7239 | 0.3845 | 0.0007 | -3.7488 | 0.0209 | -2.5583 | 0.2276 | -1.7464 |
| Precuneus R | 0.0001 | 4.0746 | 0.6862 | -0.6602 | 0.3291 | 1.1014 | 0.6006 | 0.8294 |
| Rostral anterior cingulate R | 0.7235 | -0.4009 | 0.7206 | -0.5015 | 0.0292 | -2.4194 | 0.6045 | -0.8078 |
| Rostral middle frontal R | <0.0001 | 8.8073 | 0.9310 | -0.2199 | 0.0004 | 3.7667 | 0.9631 | 0.0463 |
| Superior frontal R | 0.0013 | 3.4203 | 0.9336 | -0.1977 | 0.8987 | 0.1278 | 0.7748 | -0.4001 |
| Superior parietal R | <0.0001 | 11.7109 | 0.5364 | 0.8767 | <0.0001 | 5.4733 | 0.4684 | 1.2054 |
| Superior temporal R | <0.0001 | -4.4820 | 0.0003 | -3.9860 | <0.0001 | -5.3003 | 0.0475 | -2.6053 |
| Supramarginal R | 0.1181 | 1.7400 | 0.0694 | -2.1677 | 0.2406 | -1.3217 | 0.0589 | -2.4761 |
| Transverse temporal R | 0.0612 | 2.0490 | 0.9365 | 0.1497 | 0.8987 | 0.1495 | 0.8166 | 0.3177 |
| Insula R | <0.0001 | -4.9462 | 0.3593 | -1.2108 | <0.0001 | -7.1431 | 0.4684 | -1.1897 |

*BMI – body mass index, WHR – waist-to-hip ratio.*

**Table S6.** Associations between body mass index and waist-to-hip ratio with subcortical volume measures

| Structure | BMI |  | WHR |  |
| --- | --- | --- | --- | --- |
|  | p-value | T-value | p-value | T-value |
| Thalamus L | 0.0001 | -3.9238 | 0.0245 | -2.4152 |
| Thalamus R | 0.2218 | 1.2217 | 0.0041 | 3.0827 |
| Caudate L | 0.0002 | -3.7756 | 0.5533 | -0.5928 |
| Caudate R | 0.0156 | -2.4455 | 0.2110 | 1.2933 |
| Putamen L | <0.0001 | -8.0976 | 0.0114 | -2.7217 |
| Putamen R | <0.0001 | -13.5912 | <0.0001 | -6.3340 |
| Pallidum L | <0.0001 | -5.8545 | 0.1298 | -1.5926 |
| Pallidum R | <0.0001 | -6.2337 | 0.1298 | -1.6051 |
| Hippocampus L | <0.0001 | -5.3418 | 0.0026 | -3.2628 |
| Hippocampus R | 0.0138 | -2.5163 | 0.0517 | -2.0868 |
| Amygdala L | <0.0001 | -13.7718 | <0.0001 | -8.3091 |
| Amygdala R | <0.0001 | -13.9574 | <0.0001 | -8.5601 |
| Nucleus accumbens L | <0.0001 | -12.0100 | <0.0001 | -6.8709 |
| Nucleus accumbens R | <0.0001 | -13.9870 | <0.0001 | -7.6203 |

*BMI – body mass index, WHR – waist-to-hip ratio.*

**Table S7.** Associations between body mass index, waist-to-hip ratio and cognitive abilities.

| Cognitive ability | BMI |  | WHR |  |
| --- | --- | --- | --- | --- |
|  | p-value | estimate | p-value | estimate |
| Working memory | 0.0184 | -0.0315 | 0.0005 | -0.0594 |
| Fluid intelligence | 0.1167 | -0.0142 | 0.0001 | -0.0442 |
| Executive functions | 0.0184 | 0.0345 | 0.6496 | -0.0108 |
| Prospective memory | 0.2750 | 0.0102 | <0.0001 | 0.0540 |
| Visuospatial memory | 0.0005 | -0.0333 | 0.9346 | 0.0009 |
| Reaction time | 0.7972 | 0.0021 | 0.0011 | 0.0371 |

BMI – body mass index, WHR – waist-to-hip ratio

**Table S8.**
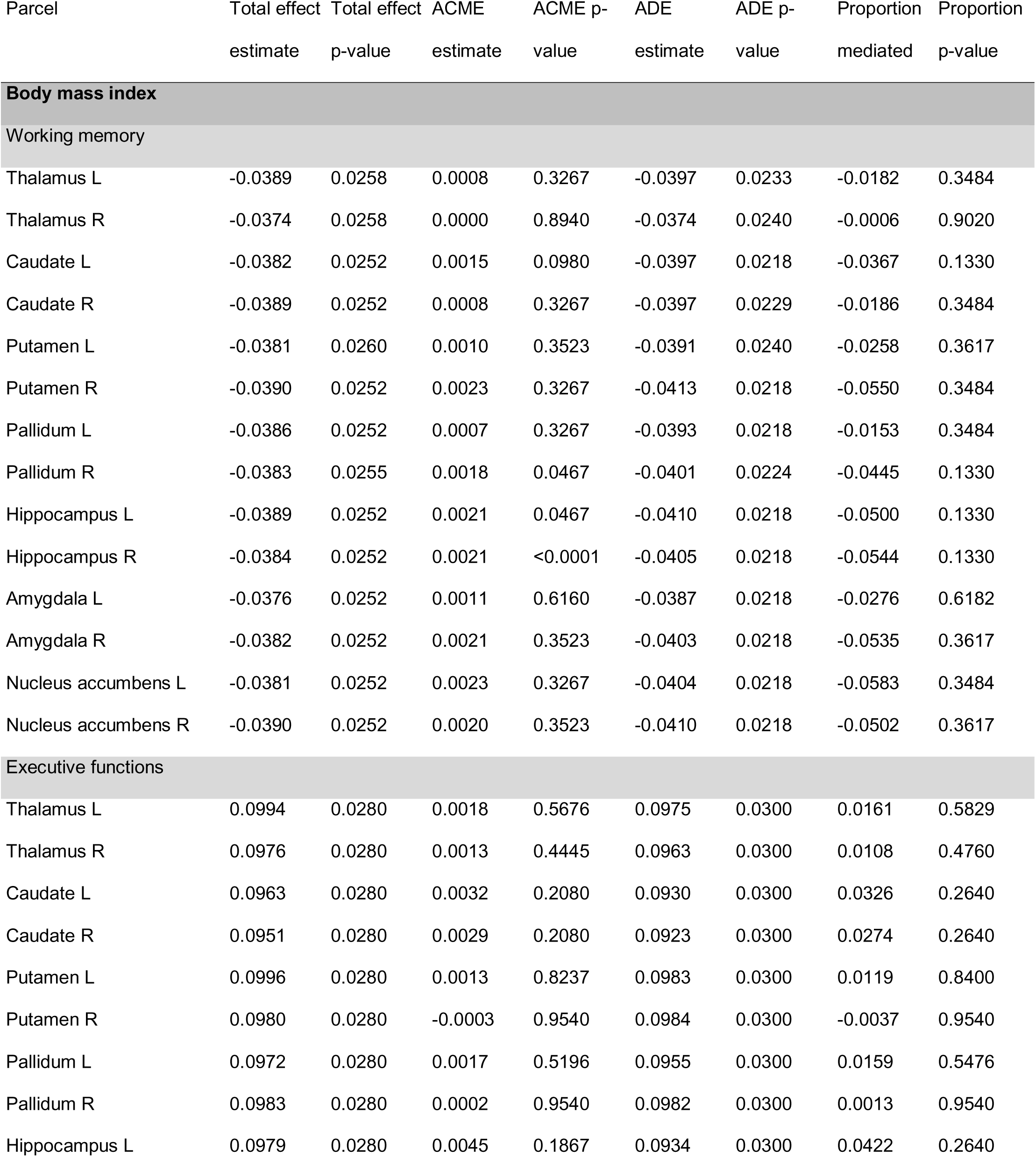
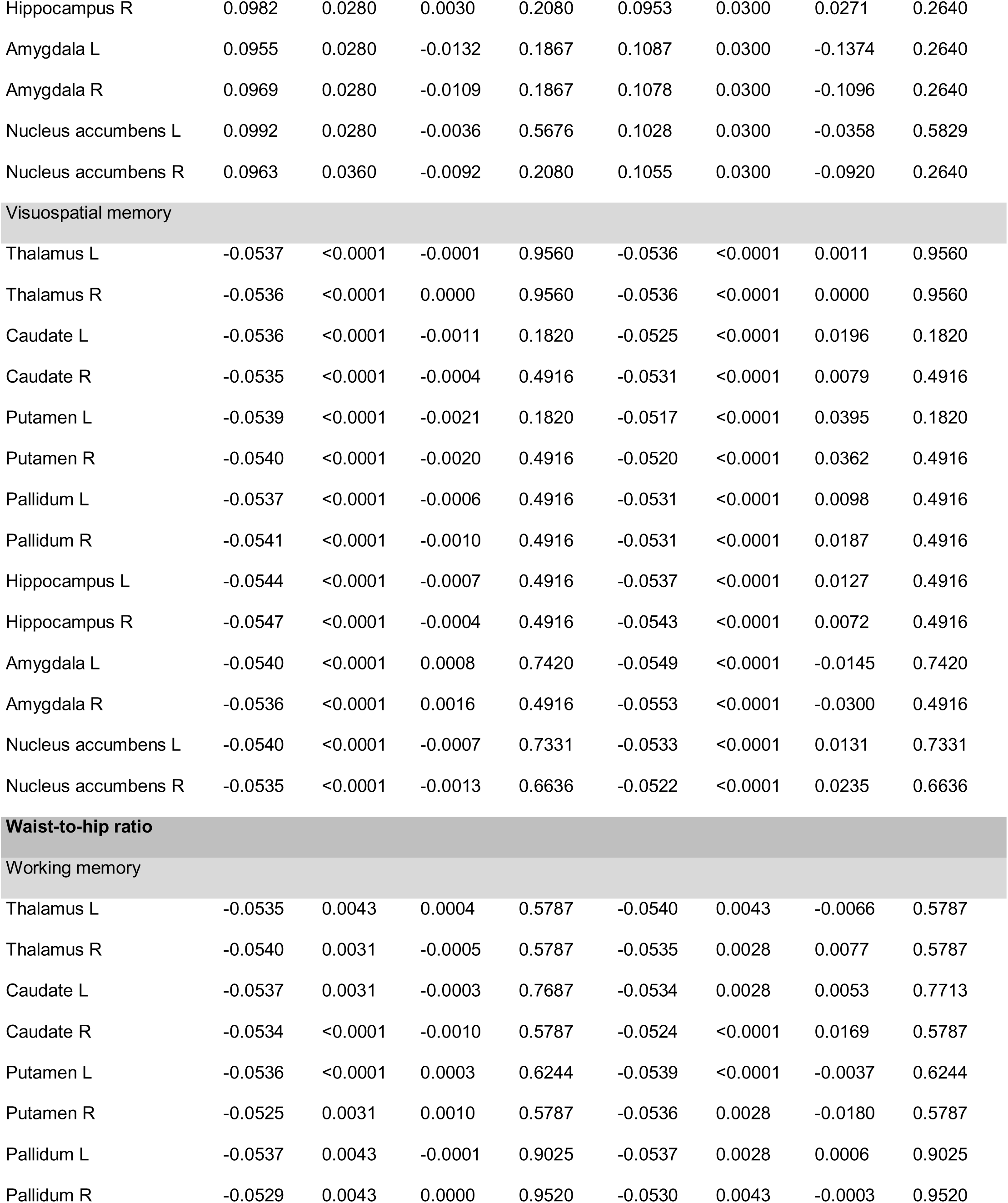
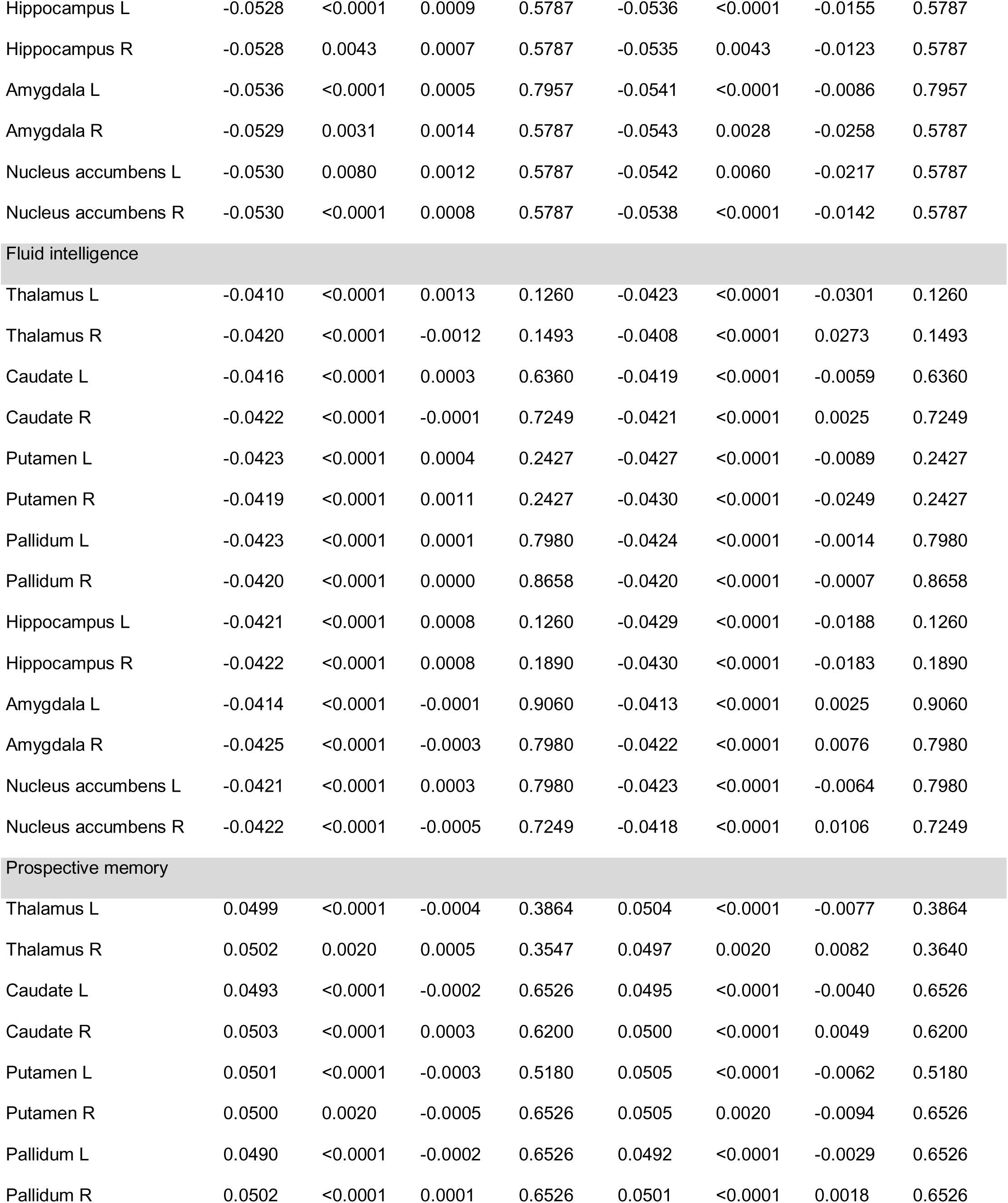
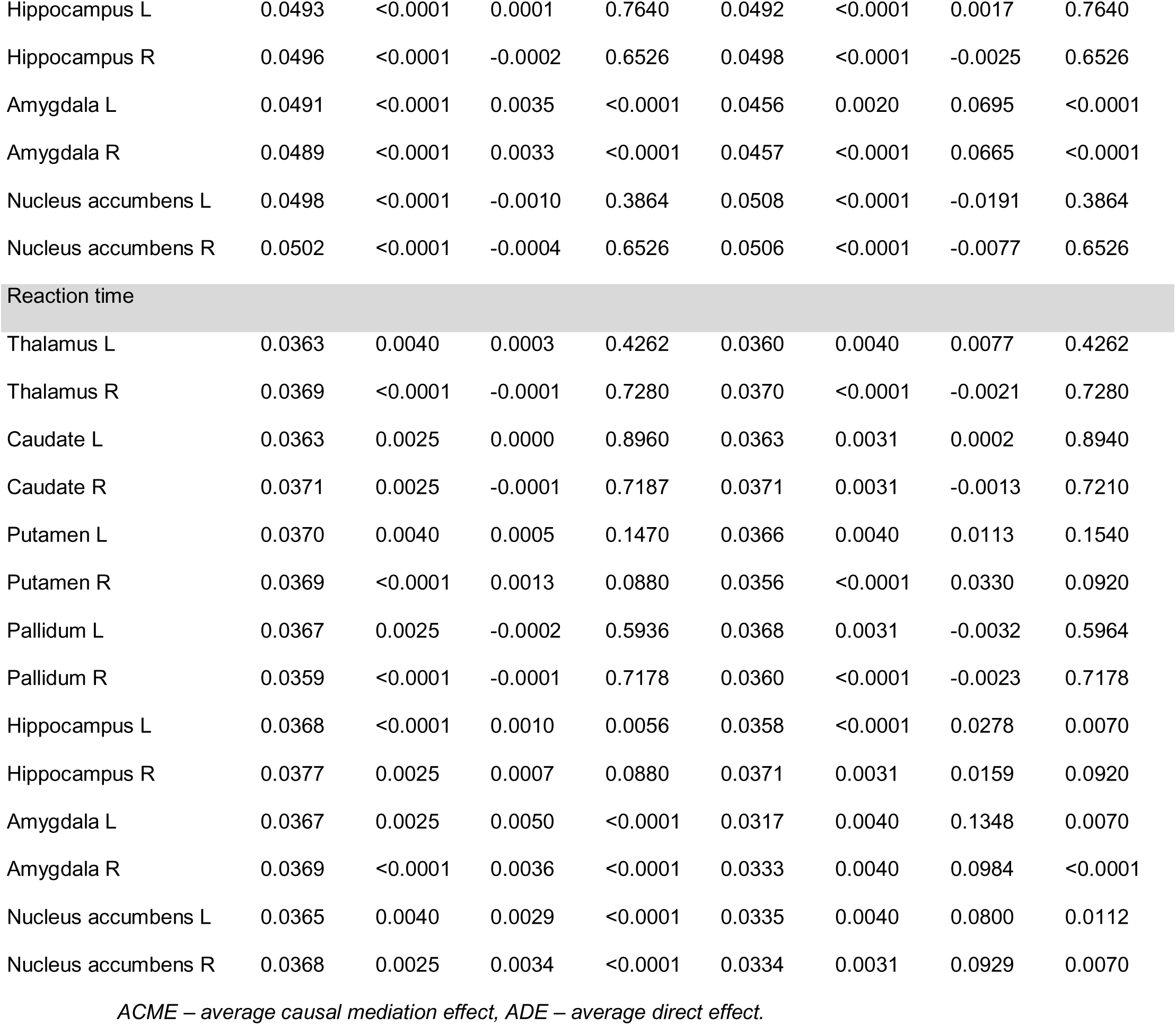
Mediation effects of subcortical grey matter volumes on the associations between body mass index and cognition, and waist-to-hip ratio and cognition.

**Table S9.** Mediation effects of white matter hyperintensities on the associations between body mass index and cortical thickness for each DKT atlas parcel.

| Parcel | Total effect estimate | Total effect p-value | ACME estimate | ACME value | p- ADE estimate | ADE value | p- Proportion mediated | Proportion p-value |
| --- | --- | --- | --- | --- | --- | --- | --- | --- |
| Caudal anterior cingulate L | -0.0357 | <0.0001 | -0.0064 | <0.0001 | -0.0293 | 0.0048 | 0.1786 | <0.0001 |
| Caudal middle frontal L | 0.0119 | 0.2237 | -0.0025 | 0.0267 | 0.0144 | 0.1153 | -0.1689 | 0.2843 |
| Cuneus L | 0.0914 | <0.0001 | -0.0009 | 0.4639 | 0.0923 | <0.0001 | -0.0101 | 0.5382 |
| Entorhinal L | -0.0563 | <0.0001 | -0.0052 | <0.0001 | -0.0511 | <0.0001 | 0.0927 | <0.0001 |
| Fusiform L | -0.0269 | 0.0038 | -0.0085 | <0.0001 | -0.0184 | 0.0541 | 0.3160 | 0.0056 |
| Inferior parietal L | 0.0152 | 0.0847 | -0.0069 | <0.0001 | 0.0221 | 0.0086 | -0.4346 | 0.1052 |
| Inferior temporal L | -0.0438 | <0.0001 | -0.0076 | <0.0001 | -0.0362 | <0.0001 | 0.1717 | <0.0001 |
| Isthmus cingulate L | -0.0037 | 0.6894 | -0.0003 | 0.7812 | -0.0034 | 0.7353 | 0.0175 | 0.9120 |
| Lateral occipital L | 0.0984 | <0.0001 | -0.0001 | 0.9120 | 0.0985 | <0.0001 | -0.0011 | 0.9120 |
| Lateral orbitofrontal L | -0.0098 | 0.3050 | -0.0097 | <0.0001 | -0.0001 | 0.9960 | 0.7828 | 0.3389 |
| Lingual L | 0.0724 | <0.0001 | 0.0049 | <0.0001 | 0.0675 | <0.0001 | 0.0677 | <0.0001 |
| Medial orbitofrontal L | -0.0181 | 0.0682 | -0.0061 | <0.0001 | -0.0119 | 0.2248 | 0.3183 | 0.0880 |
| Middle temporal L | -0.0348 | <0.0001 | -0.0069 | <0.0001 | -0.0279 | <0.0001 | 0.1982 | <0.0001 |
| Parahippocampal L | -0.0281 | <0.0001 | -0.0082 | <0.0001 | -0.0199 | 0.0086 | 0.2909 | <0.0001 |
| Paracentral L | -0.0279 | 0.0038 | -0.0154 | <0.0001 | -0.0126 | 0.1779 | 0.5437 | 0.0056 |
| Pars opercularis L | -0.0072 | 0.4492 | -0.0096 | <0.0001 | 0.0025 | 0.8017 | 0.8618 | 0.4859 |
| Pars orbitalis L | -0.0229 | 0.0073 | -0.0092 | <0.0001 | -0.0137 | 0.1522 | 0.4031 | 0.0108 |
| Pars triangularis L | 0.0171 | 0.0555 | -0.0078 | <0.0001 | 0.0249 | 0.0116 | -0.4434 | 0.0727 |
| Pericalcarine L | 0.0888 | <0.0001 | 0.0012 | 0.2591 | 0.0875 | <0.0001 | 0.0137 | 0.3297 |
| Postcentral L | 0.0531 | <0.0001 | -0.0051 | <0.0001 | 0.0581 | <0.0001 | -0.0952 | <0.0001 |
| Posterior cingulate L | -0.0106 | 0.2609 | -0.0034 | 0.0027 | -0.0072 | 0.4668 | 0.2601 | 0.3011 |
| Precentral L | -0.0091 | 0.3185 | -0.0125 | <0.0001 | 0.0034 | 0.7353 | 1.0473 | 0.3531 |
| Precuneus L | 0.0283 | <0.0001 | -0.0022 | 0.0548 | 0.0304 | <0.0001 | -0.0782 | 0.0891 |
| Rostral anterior cingulate L | -0.0340 | <0.0001 | -0.0114 | <0.0001 | -0.0226 | 0.0219 | 0.3387 | <0.0001 |
| Rostral middle frontal L | 0.0568 | <0.0001 | -0.0032 | 0.0027 | 0.0600 | <0.0001 | -0.0561 | 0.0056 |
| Superior frontal L | 0.0161 | 0.0856 | -0.0032 | 0.0027 | 0.0194 | 0.0413 | -0.1894 | 0.1094 |
| Superior parietal L | 0.0696 | <0.0001 | 0.0019 | 0.0992 | 0.0677 | <0.0001 | 0.0271 | 0.1516 |
| Superior temporal L | -0.0537 | <0.0001 | -0.0123 | <0.0001 | -0.0414 | <0.0001 | 0.2305 | <0.0001 |
| Supramarginal L | -0.0094 | 0.2834 | -0.0114 | <0.0001 | 0.0020 | 0.8638 | 0.9696 | 0.3230 |
| Transverse temporal L | 0.0051 | 0.6290 | -0.0019 | 0.0842 | 0.0070 | 0.4866 | -0.1366 | 0.6889 |
| Insula L | -0.0715 | <0.0001 | -0.0220 | <0.0001 | -0.0494 | <0.0001 | 0.3074 | <0.0001 |
| Caudal anterior cingulate R | 0.0014 | 0.8960 | -0.0059 | <0.0001 | 0.0073 | 0.4698 | -0.3615 | 0.9120 |
| Caudal middle frontal R | 0.0192 | 0.0207 | -0.0010 | 0.3851 | 0.0202 | 0.0116 | -0.0504 | 0.4702 |
| Cuneus R | 0.0891 | <0.0001 | -0.0005 | 0.6999 | 0.0896 | <0.0001 | -0.0055 | 0.7374 |
| Entorhinal R | -0.0223 | 0.0142 | -0.0068 | <0.0001 | -0.0155 | 0.1063 | 0.3010 | 0.0198 |
| Fusiform R | -0.0205 | 0.0302 | -0.0028 | 0.0127 | -0.0177 | 0.0469 | 0.1363 | 0.0620 |
| Inferior parietal R | 0.0445 | <0.0001 | -0.0078 | <0.0001 | 0.0523 | <0.0001 | -0.1754 | <0.0001 |
| Inferior temporal R | -0.0050 | 0.6085 | -0.0039 | 0.0027 | -0.0012 | 0.9197 | 0.3346 | 0.6467 |
| Isthmus cingulate R | -0.0042 | 0.6526 | -0.0042 | <0.0001 | 0.0000 | 0.9960 | 0.3641 | 0.6889 |
| Lateral occipital R | 0.0971 | <0.0001 | 0.0020 | 0.0873 | 0.0951 | <0.0001 | 0.0205 | 0.1346 |
| Lateral orbitofrontal R | 0.0133 | 0.1775 | -0.0087 | <0.0001 | 0.0219 | 0.0150 | -0.5820 | 0.2056 |
| Lingual R | 0.0771 | <0.0001 | 0.0067 | <0.0001 | 0.0704 | <0.0001 | 0.0871 | <0.0001 |
| Medial orbitofrontal R | -0.0117 | 0.2322 | -0.0057 | <0.0001 | -0.0060 | 0.5144 | 0.4181 | 0.2728 |
| Middle temporal R | 0.0045 | 0.6311 | -0.0049 | <0.0001 | 0.0094 | 0.3062 | -0.4303 | 0.6796 |
| Parahippocampal R | -0.0262 | 0.0038 | -0.0059 | <0.0001 | -0.0204 | 0.0413 | 0.2238 | 0.0056 |
| Paracentral R | -0.0276 | <0.0001 | -0.0155 | <0.0001 | -0.0122 | 0.1779 | 0.5629 | <0.0001 |
| Pars opercularis R | 0.0074 | 0.4340 | -0.0096 | <0.0001 | 0.0170 | 0.0522 | -0.8550 | 0.4702 |
| Pars orbitalis R | 0.0020 | 0.8416 | -0.0100 | <0.0001 | 0.0121 | 0.1952 | -0.7145 | 0.8851 |
| Pars triangularis R | 0.0273 | <0.0001 | -0.0072 | <0.0001 | 0.0345 | <0.0001 | -0.2656 | <0.0001 |
| Pericalcarine R | 0.0993 | <0.0001 | 0.0029 | 0.0127 | 0.0964 | <0.0001 | 0.0296 | 0.0238 |
| Postcentral R | 0.0650 | <0.0001 | -0.0056 | <0.0001 | 0.0706 | <0.0001 | -0.0861 | <0.0001 |
| Posterior cingulate R | -0.0119 | 0.2149 | -0.0043 | <0.0001 | -0.0075 | 0.4340 | 0.3222 | 0.2480 |
| Precentral R | 0.0033 | 0.6894 | -0.0108 | <0.0001 | 0.0141 | 0.0817 | -0.9968 | 0.7374 |
| Precuneus R | 0.0328 | <0.0001 | -0.0038 | <0.0001 | 0.0366 | <0.0001 | -0.1153 | <0.0001 |
| Rostral anterior cingulate R | -0.0034 | 0.7213 | -0.0083 | <0.0001 | 0.0049 | 0.6290 | 0.6876 | 0.7592 |
| Rostral middle frontal R | 0.0708 | <0.0001 | -0.0028 | 0.0174 | 0.0736 | <0.0001 | -0.0391 | 0.0321 |
| Superior frontal R | 0.0270 | <0.0001 | -0.0031 | 0.0079 | 0.0302 | <0.0001 | -0.1154 | 0.0155 |
| Superior parietal R | 0.0940 | <0.0001 | -0.0001 | 0.9120 | 0.0941 | <0.0001 | -0.0012 | 0.9120 |
| Superior temporal R | -0.0356 | <0.0001 | -0.0124 | <0.0001 | -0.0231 | 0.0086 | 0.3510 | <0.0001 |
| Supramarginal R | 0.0138 | 0.1413 | -0.0120 | <0.0001 | 0.0258 | <0.0001 | -0.8240 | 0.1642 |
| Transverse temporal R | 0.0170 | 0.0682 | -0.0064 | <0.0001 | 0.0234 | 0.0116 | -0.3582 | 0.0880 |
| Insula R | -0.0410 | <0.0001 | -0.0222 | <0.0001 | -0.0188 | 0.0558 | 0.5396 | <0.0001 |

*ACME – average causal mediation effect, ADE – average direct effect.*

**Table S10.** Mediation effects of white matter hyperintensities on the associations between body mass index and cortical volume for each DKT atlas parcel.

| Parcel | Total effect<br>estimate | Total effect<br>p-value | ACME<br>estimate | ACME<br>value | p-ADE<br>estimate | ADE<br>value | p-Proportion<br>mediated | Proportion<br>p-value |
| --- | --- | --- | --- | --- | --- | --- | --- | --- |
| Caudal anterior cingulate L | 0.0010 | 0.9493 | -0.0056 | <0.0001 | 0.0067 | 0.6464 | -0.3205 | 0.9620 |
| Caudal middle frontal L | -0.0081 | 0.4842 | -0.0024 | 0.0310 | -0.0057 | 0.6484 | 0.1875 | 0.5167 |
| Cuneus L | 0.0097 | 0.3599 | -0.0039 | <0.0001 | 0.0135 | 0.1899 | -0.3103 | 0.3689 |
| Entorhinal L | -0.0049 | 0.6735 | 0.0009 | 0.4113 | -0.0058 | 0.6464 | -0.0573 | 0.8899 |
| Fusiform L | -0.0353 | <0.0001 | -0.0079 | <0.0001 | -0.0274 | 0.0089 | 0.2240 | <0.0001 |
| Inferior parietal L | -0.0639 | <0.0001 | -0.0091 | <0.0001 | -0.0548 | <0.0001 | 0.1415 | <0.0001 |
| Inferior temporal L | -0.0506 | <0.0001 | -0.0086 | <0.0001 | -0.0419 | <0.0001 | 0.1697 | <0.0001 |
| Isthmus cingulate L | 0.0141 | 0.1356 | -0.0030 | 0.0097 | 0.0171 | 0.0809 | -0.2011 | 0.1529 |
| Lateral occipital L | 0.0100 | 0.3599 | -0.0045 | <0.0001 | 0.0145 | 0.1653 | -0.3457 | 0.3689 |
| Lateral orbitofrontal L | -0.0464 | <0.0001 | -0.0082 | <0.0001 | -0.0382 | <0.0001 | 0.1748 | <0.0001 |
| Lingual L | 0.0152 | 0.1356 | -0.0022 | 0.0513 | 0.0174 | 0.1042 | -0.1355 | 0.2090 |
| Medial orbitofrontal L | -0.0487 | <0.0001 | -0.0074 | <0.0001 | -0.0413 | <0.0001 | 0.1519 | <0.0001 |
| Middle temporal L | -0.0674 | <0.0001 | -0.0104 | <0.0001 | -0.0570 | <0.0001 | 0.1534 | <0.0001 |
| Parahippocampal L | -0.0207 | 0.0362 | -0.0058 | <0.0001 | -0.0149 | 0.1240 | 0.2764 | 0.0362 |
| Paracentral L | -0.0152 | 0.0855 | -0.0094 | <0.0001 | -0.0058 | 0.6464 | 0.6101 | 0.0855 |
| Pars opercularis L | -0.0272 | <0.0001 | -0.0065 | <0.0001 | -0.0206 | 0.0372 | 0.2403 | <0.0001 |
| Pars orbitalis L | -0.0203 | 0.0382 | -0.0058 | <0.0001 | -0.0146 | 0.1653 | 0.2834 | 0.0382 |
| Pars triangularis L | -0.0143 | 0.1422 | -0.0066 | <0.0001 | -0.0077 | 0.4627 | 0.4431 | 0.1560 |
| Pericalcarine L | 0.0049 | 0.6868 | -0.0040 | <0.0001 | 0.0089 | 0.4340 | -0.3124 | 0.7288 |
| Postcentral L | -0.0133 | 0.2032 | -0.0081 | <0.0001 | -0.0052 | 0.7027 | 0.5505 | 0.2090 |
| Posterior cingulate L | 0.0000 | 0.9620 | -0.0062 | <0.0001 | 0.0062 | 0.6464 | -0.3413 | 0.9620 |
| Precentral L | -0.0327 | 0.0065 | -0.0121 | <0.0001 | -0.0206 | 0.0219 | 0.3676 | 0.0065 |
| Precuneus L | -0.0011 | 0.9493 | -0.0027 | 0.0184 | 0.0015 | 0.9127 | 0.1407 | 0.9620 |
| Rostral anterior cingulate L | -0.0069 | 0.5676 | -0.0057 | <0.0001 | -0.0012 | 0.9127 | 0.4857 | 0.5805 |
| Rostral middle frontal L | -0.0009 | 0.9493 | -0.0028 | 0.0097 | 0.0020 | 0.8636 | 0.1470 | 0.9620 |
| Superior frontal L | 0.0021 | 0.8956 | -0.0059 | <0.0001 | 0.0080 | 0.4871 | -0.4644 | 0.9300 |
| Superior parietal L | -0.0151 | 0.1356 | -0.0026 | 0.0184 | -0.0125 | 0.2152 | 0.1660 | 0.1666 |
| Superior temporal L | -0.0344 | <0.0001 | -0.0098 | <0.0001 | -0.0245 | <0.0001 | 0.2861 | <0.0001 |
| Supramarginal L | -0.0292 | 0.0065 | -0.0066 | <0.0001 | -0.0226 | 0.0276 | 0.2277 | 0.0065 |
| Transverse temporal L | 0.0053 | 0.6735 | -0.0042 | <0.0001 | 0.0095 | 0.4079 | -0.3417 | 0.7156 |
| Insula L | -0.0257 | 0.0113 | -0.0056 | <0.0001 | -0.0201 | 0.0326 | 0.2163 | 0.0118 |
| Caudal anterior cingulate R | 0.0076 | 0.5047 | -0.0059 | <0.0001 | 0.0134 | 0.2029 | -0.5048 | 0.5167 |
| Caudal middle frontal R | -0.0053 | 0.6728 | -0.0032 | 0.0076 | -0.0021 | 0.8614 | 0.2732 | 0.6999 |
| Cuneus R | 0.0203 | 0.0382 | -0.0041 | <0.0001 | 0.0244 | 0.0089 | -0.2040 | 0.0382 |
| Entorhinal R | -0.0046 | 0.6735 | -0.0002 | 0.8820 | -0.0044 | 0.7027 | 0.0106 | 0.9620 |
| Fusiform R | -0.0346 | <0.0001 | -0.0061 | <0.0001 | -0.0285 | <0.0001 | 0.1767 | <0.0001 |
| Inferior parietal R | -0.0310 | <0.0001 | -0.0077 | <0.0001 | -0.0233 | 0.0165 | 0.2493 | <0.0001 |
| Inferior temporal R | -0.0235 | <0.0001 | -0.0054 | <0.0001 | -0.0181 | 0.0650 | 0.2300 | <0.0001 |
| Isthmus cingulate R | 0.0267 | 0.0113 | -0.0035 | 0.0052 | 0.0303 | 0.0089 | -0.1366 | 0.0216 |
| Lateral occipital R | 0.0231 | 0.0162 | -0.0046 | <0.0001 | 0.0277 | 0.0089 | -0.1995 | 0.0169 |
| Lateral orbitofrontal R | -0.0474 | <0.0001 | -0.0085 | <0.0001 | -0.0390 | <0.0001 | 0.1786 | <0.0001 |
| Lingual R | 0.0139 | 0.1771 | -0.0002 | 0.8820 | 0.0141 | 0.1680 | -0.0102 | 0.9620 |
| Medial orbitofrontal R | -0.0377 | <0.0001 | -0.0094 | <0.0001 | -0.0283 | <0.0001 | 0.2478 | <0.0001 |
| Middle temporal R | -0.0366 | <0.0001 | -0.0085 | <0.0001 | -0.0281 | <0.0001 | 0.2348 | <0.0001 |
| Parahippocampal R | -0.0216 | 0.0113 | -0.0049 | <0.0001 | -0.0166 | 0.0982 | 0.2326 | 0.0118 |
| Paracentral R | -0.0103 | 0.3402 | -0.0091 | <0.0001 | -0.0012 | 0.9127 | 0.7272 | 0.3492 |
| Pars opercularis R | -0.0119 | 0.2514 | -0.0073 | <0.0001 | -0.0046 | 0.7027 | 0.5331 | 0.2583 |
| Pars orbitalis R | -0.0222 | 0.0065 | -0.0072 | <0.0001 | -0.0151 | 0.1506 | 0.3235 | 0.0065 |
| Pars triangularis R | -0.0048 | 0.6735 | -0.0058 | <0.0001 | 0.0011 | 0.9127 | 0.5255 | 0.7156 |
| Pericalcarine R | 0.0148 | 0.1422 | -0.0023 | 0.0348 | 0.0171 | 0.1240 | -0.1454 | 0.2029 |
| Postcentral R | 0.0010 | 0.9493 | -0.0080 | <0.0001 | 0.0090 | 0.4340 | -0.5988 | 0.9620 |
| Posterior cingulate R | -0.0176 | 0.0753 | -0.0060 | <0.0001 | -0.0116 | 0.3189 | 0.3331 | 0.0753 |
| Precentral R | -0.0311 | 0.0065 | -0.0117 | <0.0001 | -0.0193 | 0.0676 | 0.3767 | 0.0065 |
| Precuneus R | -0.0056 | 0.6728 | -0.0026 | 0.0167 | -0.0030 | 0.8496 | 0.2050 | 0.7009 |
| Rostral anterior cingulate R | -0.0040 | 0.7510 | -0.0062 | <0.0001 | 0.0022 | 0.8614 | 0.5356 | 0.7961 |
| Rostral middle frontal R | -0.0017 | 0.9493 | -0.0019 | 0.0694 | 0.0002 | 0.9320 | 0.0861 | 0.9620 |
| Superior frontal R | -0.0015 | 0.9493 | -0.0050 | <0.0001 | 0.0035 | 0.7885 | 0.3467 | 0.9620 |
| Superior parietal R | 0.0074 | 0.5495 | -0.0027 | 0.0271 | 0.0101 | 0.3548 | -0.2125 | 0.5805 |
| Superior temporal R | -0.0326 | <0.0001 | -0.0095 | <0.0001 | -0.0231 | 0.0219 | 0.2961 | <0.0001 |
| Supramarginal R | -0.0181 | 0.0505 | -0.0082 | <0.0001 | -0.0098 | 0.4079 | 0.4505 | 0.0505 |
| Transverse temporal R | 0.0008 | 0.9493 | -0.0059 | <0.0001 | 0.0067 | 0.5912 | -0.3463 | 0.9620 |
| Insula R | -0.0107 | 0.3231 | -0.0065 | <0.0001 | -0.0041 | 0.7535 | 0.5122 | 0.3318 |

*ACME – average causal mediation effect, ADE – average direct effect.*

**Table S11.** Mediation effects of white matter hyperintensities on the associations between waist-to-hip ratio and cortical thickness for each DKT atlas parcel.

| Parcel | Total effect estimate | Total effect p-value | ACME estimate | ACME value | p- ADE estimate | ADE value | p- Proportion mediated | Proportion p-value |
| --- | --- | --- | --- | --- | --- | --- | --- | --- |
| Caudal anterior cingulate L | -0.0479 | <0.0001 | -0.0091 | <0.0001 | -0.0388 | 0.0054 | 0.1876 | <0.0001 |
| Caudal middle frontal L | -0.0153 | 0.2111 | -0.0030 | 0.0668 | -0.0124 | 0.3467 | 0.1634 | 0.2875 |
| Cuneus L | 0.0665 | <0.0001 | -0.0003 | 0.8619 | 0.0668 | <0.0001 | -0.0053 | 0.9129 |
| Entorhinal L | -0.0686 | <0.0001 | -0.0074 | <0.0001 | -0.0612 | <0.0001 | 0.1086 | <0.0001 |
| Fusiform L | -0.0624 | <0.0001 | -0.0114 | <0.0001 | -0.0510 | <0.0001 | 0.1850 | <0.0001 |
| Inferior parietal L | 0.0056 | 0.6264 | -0.0094 | <0.0001 | 0.0150 | 0.2450 | -0.6403 | 0.6606 |
| Inferior temporal L | -0.0728 | <0.0001 | -0.0104 | <0.0001 | -0.0624 | <0.0001 | 0.1417 | <0.0001 |
| Isthmus cingulate L | -0.0138 | 0.2951 | -0.0003 | 0.8619 | -0.0135 | 0.3245 | 0.0182 | 0.9129 |
| Lateral occipital L | 0.0708 | <0.0001 | 0.0009 | 0.6011 | 0.0699 | <0.0001 | 0.0123 | 0.6606 |
| Lateral orbitofrontal L | -0.0208 | 0.1096 | -0.0136 | <0.0001 | -0.0073 | 0.5558 | 0.6210 | 0.1240 |
| Lingual L | 0.0601 | <0.0001 | 0.0076 | <0.0001 | 0.0525 | <0.0001 | 0.1259 | <0.0001 |
| Medial orbitofrontal L | -0.0416 | <0.0001 | -0.0083 | <0.0001 | -0.0334 | 0.0103 | 0.1972 | <0.0001 |
| Middle temporal L | -0.0564 | <0.0001 | -0.0095 | <0.0001 | -0.0469 | <0.0001 | 0.1687 | <0.0001 |
| Parahippocampal L | -0.0303 | <0.0001 | -0.0117 | <0.0001 | -0.0186 | 0.1616 | 0.3883 | <0.0001 |
| Paracentral L | -0.0438 | <0.0001 | -0.0216 | <0.0001 | -0.0222 | 0.0964 | 0.4880 | <0.0001 |
| Pars opercularis L | -0.0276 | 0.0165 | -0.0133 | <0.0001 | -0.0144 | 0.2624 | 0.4812 | 0.0225 |
| Pars orbitalis L | -0.0218 | 0.0858 | -0.0132 | <0.0001 | -0.0086 | 0.5231 | 0.5885 | 0.1015 |
| Pars triangularis L | -0.0068 | 0.6156 | -0.0105 | <0.0001 | 0.0037 | 0.7419 | 0.6795 | 0.6606 |
| Pericalcarine L | 0.0773 | <0.0001 | 0.0025 | 0.1174 | 0.0748 | <0.0001 | 0.0316 | 0.1643 |
| Postcentral L | 0.0292 | 0.0225 | -0.0064 | <0.0001 | 0.0357 | 0.0054 | -0.2188 | 0.0298 |
| Posterior cingulate L | -0.0250 | 0.0496 | -0.0046 | 0.0081 | -0.0205 | 0.1447 | 0.1784 | 0.0680 |
| Precentral L | -0.0378 | <0.0001 | -0.0171 | <0.0001 | -0.0207 | 0.1326 | 0.4524 | <0.0001 |
| Precuneus L | 0.0074 | 0.5614 | -0.0025 | 0.1171 | 0.0099 | 0.4547 | -0.1438 | 0.6606 |
| Rostral anterior cingulate L | -0.0516 | <0.0001 | -0.0160 | <0.0001 | -0.0356 | <0.0001 | 0.3135 | <0.0001 |
| Rostral middle frontal L | 0.0157 | 0.2111 | -0.0034 | 0.0267 | 0.0191 | 0.2019 | -0.1865 | 0.2567 |
| Superior frontal L | -0.0178 | 0.1127 | -0.0038 | 0.0081 | -0.0140 | 0.2818 | 0.2020 | 0.1367 |
| Superior parietal L | 0.0433 | <0.0001 | 0.0036 | 0.0267 | 0.0397 | <0.0001 | 0.0818 | 0.0487 |
| Superior temporal L | -0.0839 | <0.0001 | -0.0171 | <0.0001 | -0.0668 | <0.0001 | 0.2036 | <0.0001 |
| Supramarginal L | -0.0330 | <0.0001 | -0.0157 | <0.0001 | -0.0173 | 0.1787 | 0.4724 | <0.0001 |
| Transverse temporal L | -0.0080 | 0.5614 | -0.0024 | 0.1174 | -0.0055 | 0.6620 | 0.1377 | 0.6606 |
| Insula L | -0.1042 | <0.0001 | -0.0309 | <0.0001 | -0.0733 | <0.0001 | 0.2949 | <0.0001 |
| Caudal anterior cingulate R | -0.0177 | 0.1598 | -0.0079 | <0.0001 | -0.0098 | 0.4547 | 0.4060 | 0.1754 |
| Caudal middle frontal R | -0.0013 | 0.9220 | -0.0009 | 0.5837 | -0.0004 | 0.9800 | 0.0174 | 0.9160 |
| Cuneus R | 0.0892 | <0.0001 | -0.0003 | 0.8960 | 0.0895 | <0.0001 | -0.0023 | 0.9148 |
| Entorhinal R | -0.0227 | 0.0816 | -0.0097 | <0.0001 | -0.0131 | 0.3436 | 0.4111 | 0.0969 |
| Fusiform R | -0.0277 | 0.0225 | -0.0040 | 0.0129 | -0.0237 | 0.0858 | 0.1448 | 0.0487 |
| Inferior parietal R | 0.0134 | 0.2784 | -0.0102 | <0.0001 | 0.0236 | 0.0645 | -0.6039 | 0.3031 |
| Inferior temporal R | -0.0115 | 0.3529 | -0.0054 | 0.0029 | -0.0062 | 0.6331 | 0.3282 | 0.3931 |
| Isthmus cingulate R | -0.0205 | 0.0938 | -0.0056 | <0.0001 | -0.0149 | 0.2624 | 0.2626 | 0.1098 |
| Lateral occipital R | 0.0856 | <0.0001 | 0.0036 | 0.0129 | 0.0820 | <0.0001 | 0.0424 | 0.0270 |
| Lateral orbitofrontal R | 0.0100 | 0.4141 | -0.0121 | <0.0001 | 0.0221 | 0.1151 | -0.7895 | 0.4573 |
| Lingual R | 0.0756 | <0.0001 | 0.0099 | <0.0001 | 0.0657 | <0.0001 | 0.1302 | <0.0001 |
| Medial orbitofrontal R | -0.0221 | 0.0974 | -0.0079 | <0.0001 | -0.0142 | 0.2948 | 0.3315 | 0.1106 |
| Middle temporal R | -0.0209 | 0.0899 | -0.0065 | <0.0001 | -0.0145 | 0.2450 | 0.3032 | 0.1058 |
| Parahippocampal R | -0.0260 | 0.0401 | -0.0085 | <0.0001 | -0.0175 | 0.2044 | 0.3195 | 0.0487 |
| Paracentral R | -0.0324 | <0.0001 | -0.0219 | <0.0001 | -0.0105 | 0.4304 | 0.6846 | <0.0001 |
| Pars opercularis R | -0.0237 | 0.0503 | -0.0130 | <0.0001 | -0.0108 | 0.4216 | 0.5324 | 0.0620 |
| Pars orbitalis R | 0.0058 | 0.6264 | -0.0143 | <0.0001 | 0.0201 | 0.1616 | -0.9576 | 0.6606 |
| Pars triangularis R | 0.0056 | 0.6473 | -0.0097 | <0.0001 | 0.0152 | 0.2447 | -0.6161 | 0.6820 |
| Pericalcarine R | 0.1027 | <0.0001 | 0.0046 | 0.0081 | 0.0981 | <0.0001 | 0.0449 | 0.0177 |
| Postcentral R | 0.0343 | 0.0043 | -0.0070 | <0.0001 | 0.0413 | <0.0001 | -0.2008 | 0.0062 |
| Posterior cingulate R | -0.0241 | 0.0503 | -0.0060 | <0.0001 | -0.0181 | 0.1616 | 0.2464 | 0.0620 |
| Precentral R | -0.0271 | <0.0001 | -0.0147 | <0.0001 | -0.0124 | 0.3181 | 0.5437 | <0.0001 |
| Precuneus R | 0.0120 | 0.3307 | -0.0048 | 0.0029 | 0.0168 | 0.2044 | -0.2936 | 0.3693 |
| Rostral anterior cingulate R | -0.0271 | 0.0225 | -0.0113 | <0.0001 | -0.0158 | 0.2448 | 0.4164 | 0.0298 |
| Rostral middle frontal R | 0.0411 | <0.0001 | -0.0030 | 0.0749 | 0.0441 | <0.0001 | -0.0726 | 0.1102 |
| Superior frontal R | 0.0017 | 0.9114 | -0.0038 | 0.0177 | 0.0055 | 0.6521 | -0.1611 | 0.9148 |
| Superior parietal R | 0.0589 | <0.0001 | 0.0011 | 0.5308 | 0.0578 | <0.0001 | 0.0175 | 0.6175 |
| Superior temporal R | -0.0565 | <0.0001 | -0.0174 | <0.0001 | -0.0391 | <0.0001 | 0.3100 | <0.0001 |
| Supramarginal R | -0.0143 | 0.2273 | -0.0164 | <0.0001 | 0.0021 | 0.8761 | 0.9732 | 0.2567 |
| Transverse temporal R | 0.0013 | 0.9148 | -0.0087 | <0.0001 | 0.0100 | 0.4547 | -0.4093 | 0.9148 |
| Insula R | -0.0794 | <0.0001 | -0.0309 | <0.0001 | -0.0485 | <0.0001 | 0.3867 | <0.0001 |

*ACME – average causal mediation effect, ADE – average direct effect.*

**Table S12.**
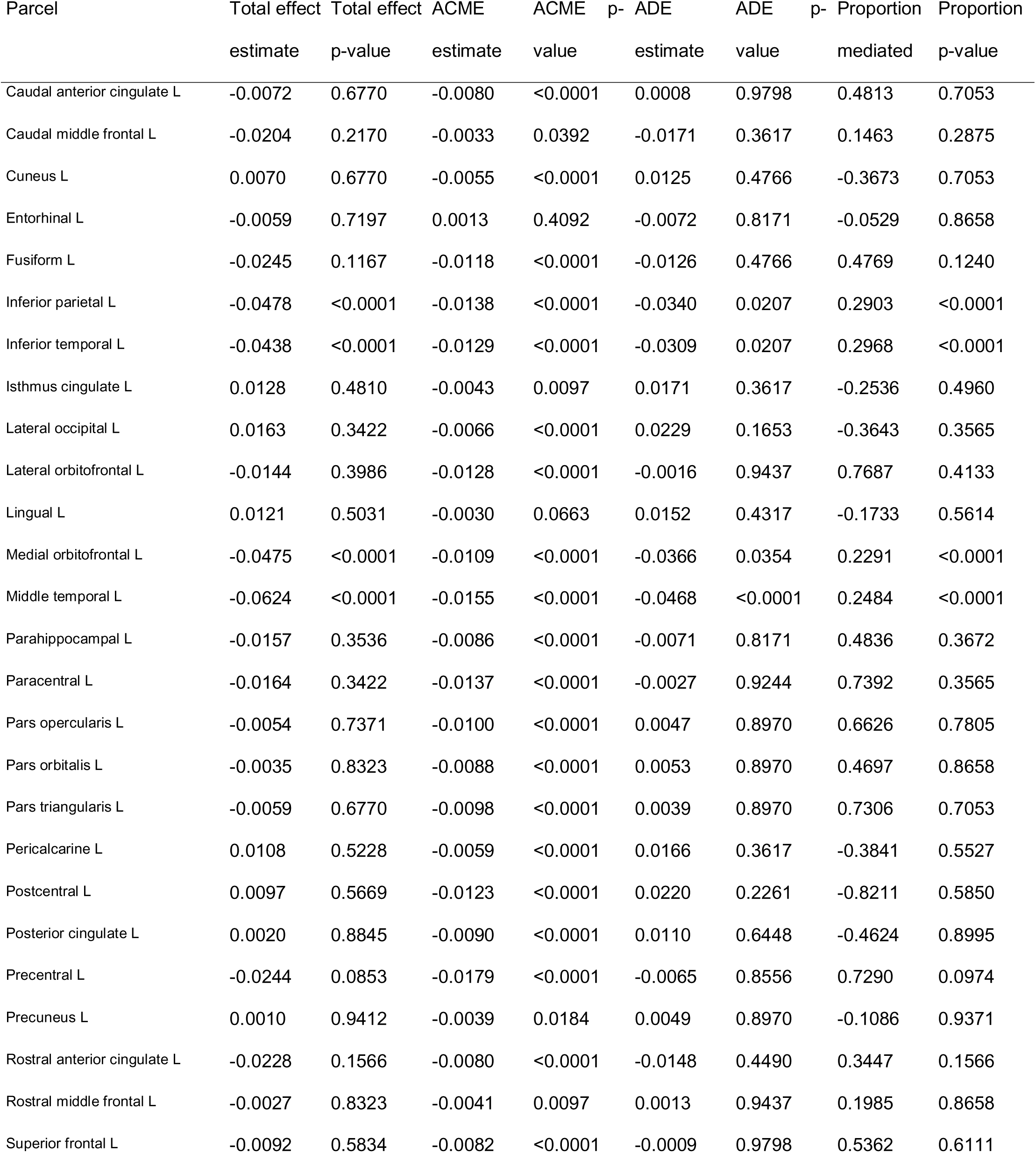
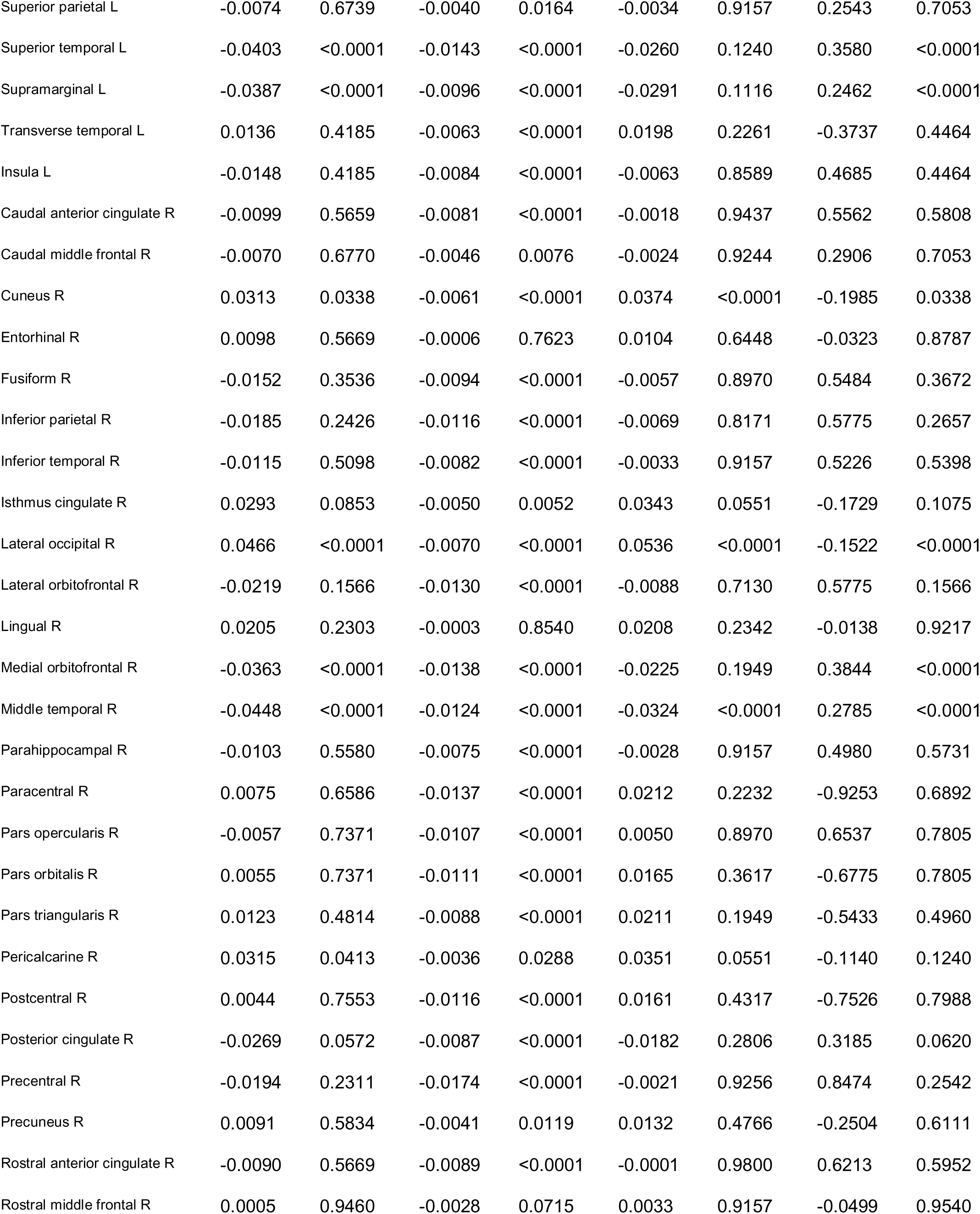
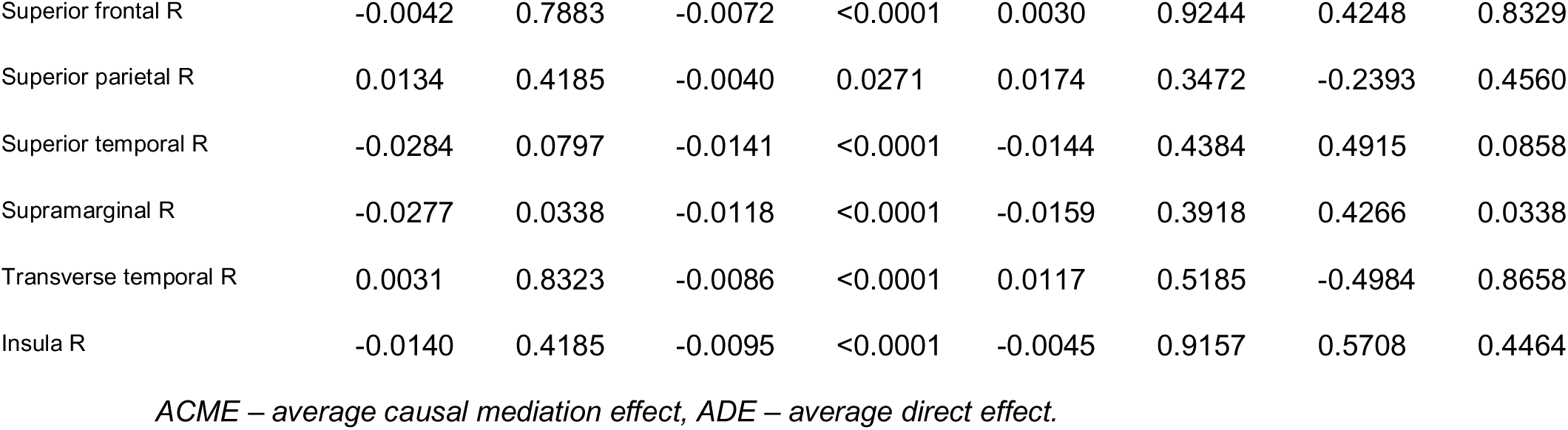
Mediation effects of white matter hyperintensities on the associations between waist-to-hip ratio and cortical volume for each DKT atlas parcel.

**Table S13.**
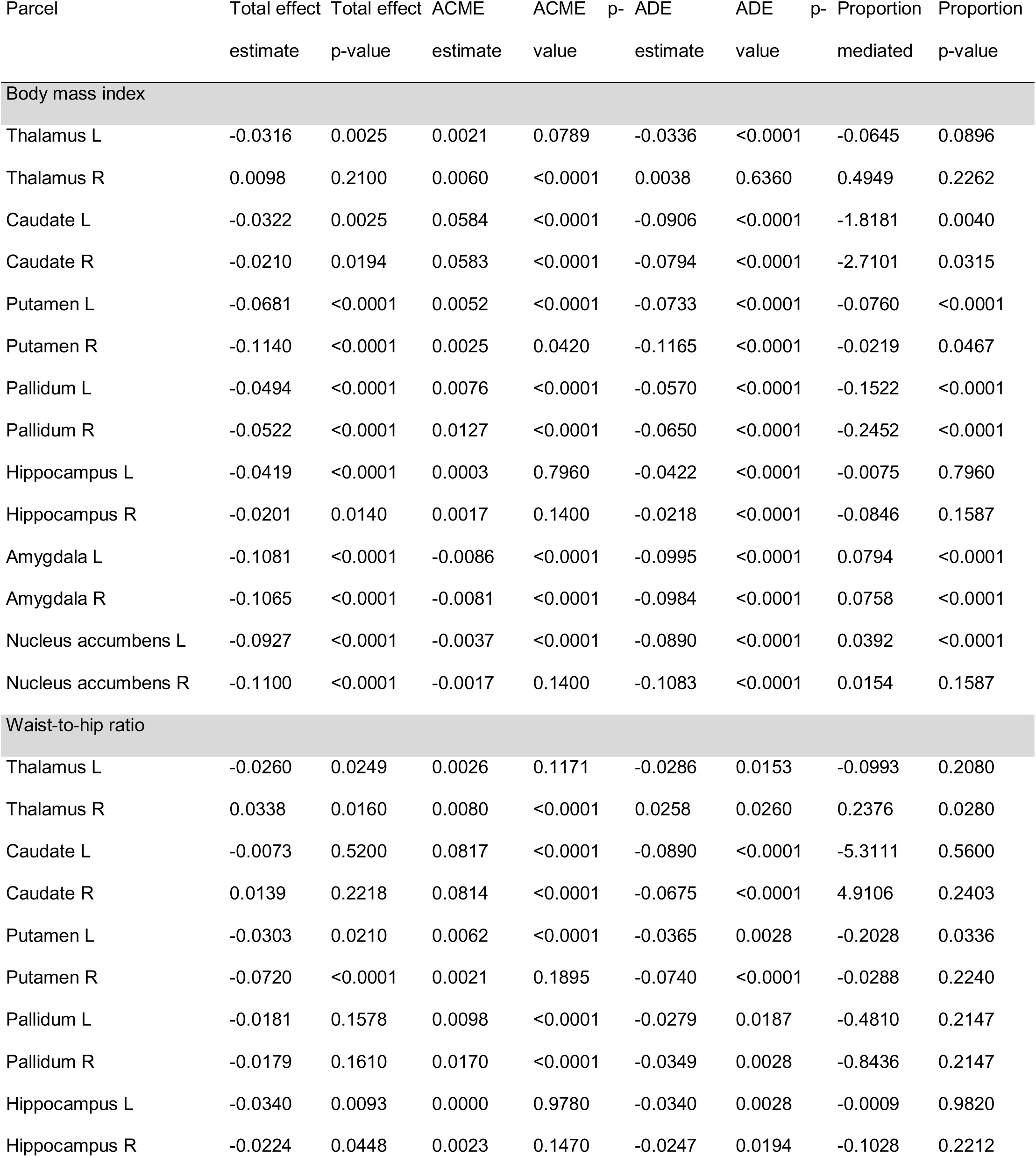
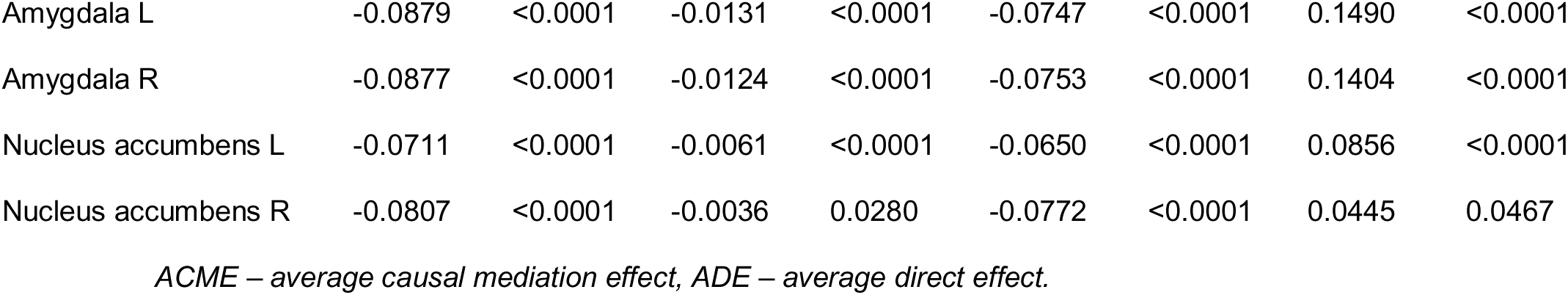
Mediation effects of white matter hyperintensities on the associations between body mass index and subcortical volumes, and waist-to-hip ratio and subcortical volumes.

**Table S14.**
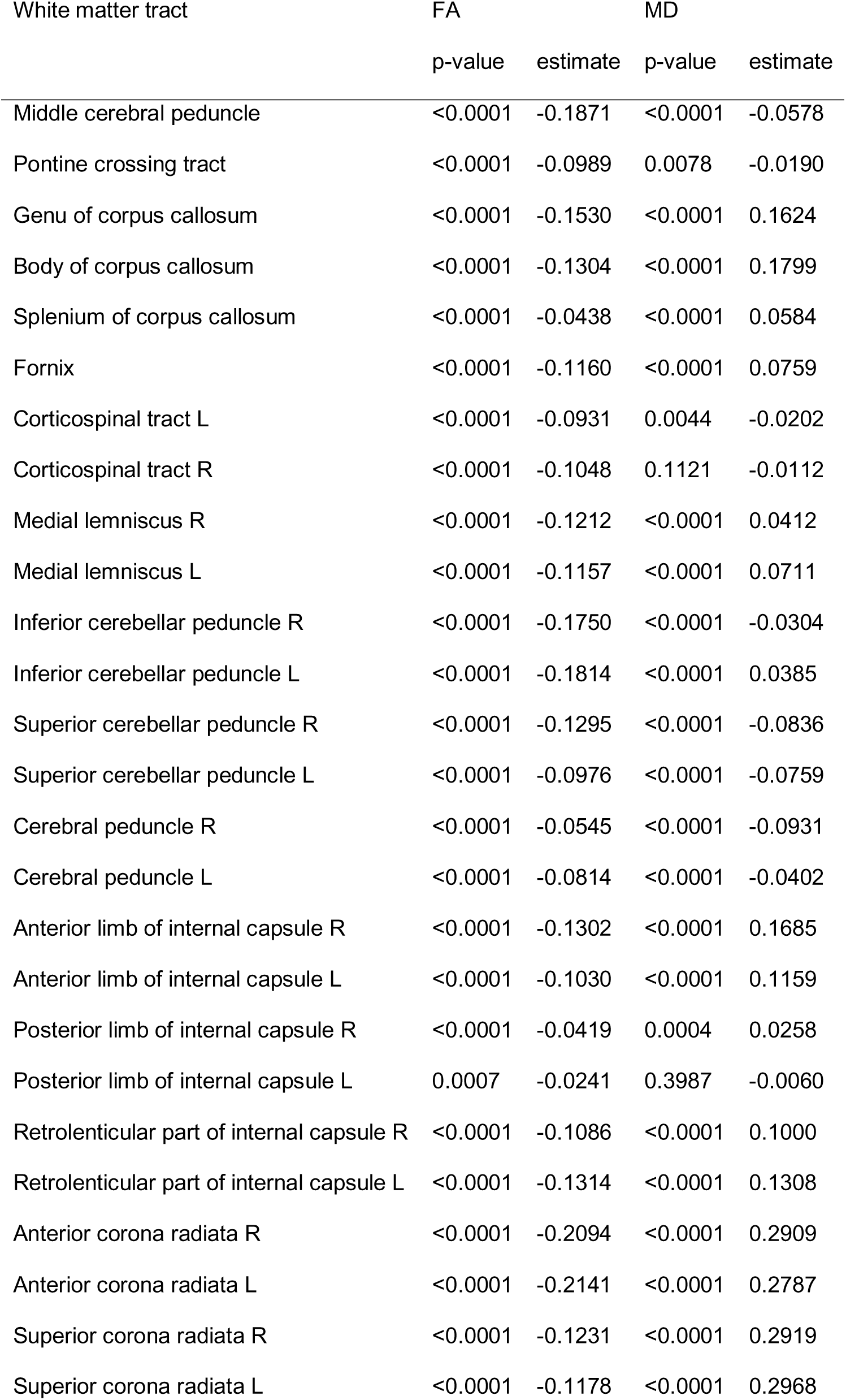
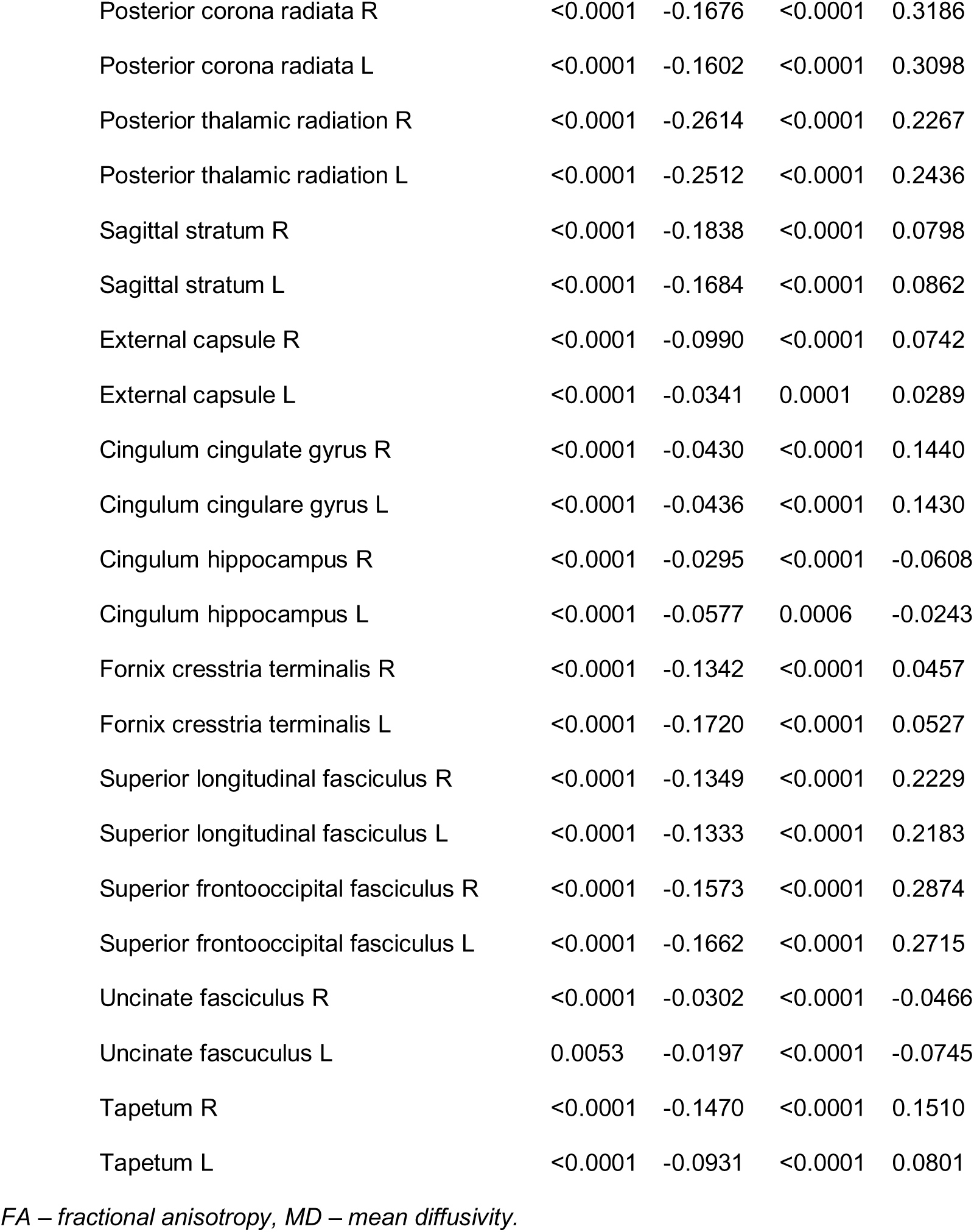
Associations between white matter hyperintensities, fractional anisotropy and mean diffusivity.

## References

Abraham, A., Pedregosa, F., Eickenberg, M., Gervais, P., Mueller, A., Kossaifi, J., Gramfort, A., Thirion, B., and Varoquaux, G. (2014). Machine learning for neuroimaging with scikit-learn. Front. Neuroinform. 8, 14.

Alarcón, G., Ray, S., and Nagel, B.J. (2016). Lower Working Memory Performance in Overweight and Obese Adolescents Is Mediated by White Matter Microstructure. J. Int. Neuropsychol. Soc. 22, 281–292.

Alba-Ferrara, L.M., and de Erausquin, G.A. (2013). What does anisotropy measure? Insights from increased and decreased anisotropy in selective fiber tracts in schizophrenia. Front. Integr. Neurosci. 7, 9.

Alexander, A.L., Lee, J.E., Lazar, M., and Field, A.S. (2007). Diffusion Tensor Imaging of the Brain. Neurotherapeutics 4, 316–329.

Alfaro-Almagro, F., Jenkinson, M., Bangerter, N.K., Andersson, J.L.R., Griffanti, L., Douaud, G., Sotiropoulos, S.N., Jbabdi, S., Hernandez-Fernandez, M., Vallee, E., et al. (2018). Image processing and Quality Control for the first 10,000 brain imaging datasets from UK Biobank. Neuroimage 166, 400–424.

Alford, S., Patel, D., Perakakis, N., and Mantzoros, C.S. (2018). Obesity as a risk factor for Alzheimer’s disease: weighing the evidence. Obes. Rev. 19, 269–280.

Apovian, C.M. (2016). Obesity: definition, comorbidities, causes, and burden. Am. J. Manag. Care 22, s176–185.

Aribisala, B.S., Valdés Hernández, M.C., Royle, N.A., Morris, Z., Muñoz Maniega, S., Bastin, M.E., Deary, I.J., and Wardlaw, J.M. (2013). Brain atrophy associations with white matter lesions in the ageing brain: The Lothian Birth Cohort 1936. Eur. Radiol. 23, 1084–1092.

Bailey, E.L., Smith, C., Sudlow, C.L.M., and Wardlaw, J.M. (2012). Pathology of lacunar ischemic stroke in humans - A systematic review. Brain Pathol. 22, 583–591.

Baron, R.M., and Kenny, D.A. (1986). The moderator-mediator variable distinction in social psychological research: Conceptual, strategic, and statistical considerations. J. Pers. Soc. Psychol. 51, 1173–1182.

Basser, P.J. (1995). Inferring microstructural features and the physiological state of tissues from diffusion□weighted images. NMR Biomed. 8, 333–344.

Bendlin, B.B., Fitzgerald, M.E., Ries, M.L., Xu, G., Kastman, E.K., Thiel, B.W., Rowley, H.A., Lazar, M., Alexander, A.L., and Johnson, S.C. (2010). White matter in aging and cognition: A cross-sectional study of microstructure in adults aged eighteen to eighty-three. Dev. Neuropsychol. 35, 257–277.

Benito-León, J., Mato-Abad, V., Louis, E.D., Hernández-Tamames, J.A., Álvarez-Linera, J., Bermejo-Pareja, F., Domingo-Santos, Á., Collado, L., and Romero, J.P. (2017). White matter microstructural changes are related to cognitive dysfunction in essential tremor. Sci. Rep. 7, 1–11.

Benjamini, Y., and Hochberg, Y. (1995). Controlling the False Discovery Rate: A Practical and Powerful Approach to Multiple Testing. J. R. Stat. Soc. Ser. B 57, 289–300.

Beyer, F., Masouleh, S.K., Kratzsch, J., Schroeter, M.L., Röhr, S., Riedel-Heller, S.G., Villringer, A., and Veronica Witte, A. (2019a). A metabolic obesity profile is associated with decreased gray matter volume in cognitively healthy older adults. Front. Aging Neurosci. 10, 202.

Beyer, F., García-García, I., Heinrich, M., Schroeter, M.L., Sacher, J., Luck, T., Riedel-Heller, S.G., Stumvoll, M., Villringer, A., and Witte, A.V. (2019b). Neuroanatomical correlates of food addiction symptoms and body mass index in the general population. Hum. Brain Mapp. 1–12.

Bowman, G.L., Kaye, J.A., and Quinn, J.F. (2012). Dyslipidemia and Blood-Brain Barrier Integrity in Alzheimer’s Disease. Curr. Gerontol. Geriatr. Res. 2012, 184042.

Castellon, X., and Bogdanova, V. (2016). Chronic inflammatory diseases and endothelial dysfunction. Aging Dis. 7, 81.

Chard, D.T., Parker, G.J.M., Griffin, C.M.B., Thompson, A.J., and Miller, D.H. (2002). The reproducibility and sensitivity of brain tissue volume measurements derived from an SPM-based segmentation methodology. J. Magn. Reson. Imaging 15, 259–267.

Chard, D.T., Jackson, J.S., Miller, D.H., and Wheeler-Kingshott, C.A.M. (2010). Reducing the impact of white matter lesions on automated measures of brain gray and white matter volumes. J. Magn. Reson. Imaging 32, 223–228.

Cullen, B., Nicholl, B.I., Mackay, D.F., Martin, D., Ul-Haq, Z., McIntosh, A., Gallacher, J., Deary, I.J., Pell, J.P., Evans, J.J., et al. (2015). Cognitive function and lifetime features of depression and bipolar disorder in a large population sample: Cross-sectional study of 143,828 UK Biobank participants. Eur. Psychiatry 30, 950–958.

Cusi, K. (2010). The role of adipose tissue and lipotoxicity in the pathogenesis of type 2 diabetes. Curr. Diab. Rep. 10, 306–315.

Dadar, M., Zeighami, Y., Yau, Y., Fereshtehnejad, S.M., Maranzano, J., Postuma, R.B., Dagher, A., and Collins, D.L. (2018). White matter hyperintensities are linked to future cognitive decline in de novo Parkinson’s disease patients. NeuroImage Clin. 20, 892–900.

Dadar, M., Potvin, O., Richard, C., and Duchesne, S. (2020). White Matter Hyperintensities Cause Systematic Errors in FreeSurfer Grey Matter Segmentations. Manuscr. Prep.

Debette, S., and Markus, H.S. (2010). The clinical importance of white matter hyperintensities on brain magnetic resonance imaging: Systematic review and meta-analysis. BMJ 341, 288.

Debette, S., Beiser, A., Hoffmann, U., DeCarli, C., O’Donnell, C.J., Massaro, J.M., Au, R., Himali, J.J., Wolf, P.A., Fox, C.S., et al. (2010). Visceral fat is associated with lower brain volume in healthy middle-aged adults. Ann. Neurol. 68, 136–144.

Dekkers, I.A., Jansen, P.R., and Lamb, H.J. (2019). Obesity, Brain Volume, and White Matter Microstructure at MRI: A Cross-sectional UK Biobank Study. Radiology 181012.

Van Dijk, E.J., Prins, N.D., Vermeer, S.E., Vrooman, H.A., Hofman, A., Koudstaal, P.J., and Breteler, M.M.B. (2005). C-reactive protein and cerebral small-vessel disease: The Rotterdam scan study. Circulation 112, 900–905.

DiSabato, D.J., Quan, N., and Godbout, J.P. (2016). Neuroinflammation: the devil is in the details. J. Neurochem. 139, 136–153.

Dobbelsteyn, C.J., Joffres, M.R., MacLean, D.R., Flowerdew, G., Balram, C., Blair, L., Butler-Jones, D., Cameron, R., Collins-Nakai, R., Connelly, P.W., et al. (2001). A comparative evaluation of waist circumference, waist-to-hip ratio and body mass index as indicators of cardiovascular risk factors. The Canadian heart health surveys. Int. J. Obes. 25, 652–661.

Dufouil, C., De Kersaint-Gilly, A., Besançon, V., Levy, C., Auffray, E., Brunnereau, L., Alpérovitch, A., and Tzourio, C. (2001). Longitudinal study of blood pressure and white matter hyperintensities: The EVA MRI cohort. Neurology 56, 921–926.

Dufouil, C., Chalmers, J., Coskun, O., Besançon, V., Bousser, M.G., Guillon, P., MacMahon, S., Mazoyer, B., Neal, B., Woodward, M., et al. (2005). Effects of blood pressure lowering on cerebral white matter hyperintensities in patients with stroke: The PROGRESS (Perindopril Protection Against Recurrent Stroke Study) Magnetic Resonance Imaging Substudy. Circulation 112, 1644–1650.

Elliott, P., Peakman, T.C., and UK Biobank (2008). The UK Biobank sample handling and storage protocol for the collection, processing and archiving of human blood and urine. Int. J. Epidemiol. 37, 234–244.

Fawns-Ritchie, C., and Deary, I.J. (2019). Reliability and validity of the UK Biobank cognitive tests. MedRxiv 19002204.

Fischl, B. (2012). FreeSurfer. Neuroimage 62, 774–781.

Fjell, A.M., Walhovd, K.B., Fennema-Notestine, C., McEvoy, L.K., Hagler, D.J., Holland, D., Brewer, J.B., and Dale, A.M. (2009). One-year brain atrophy evident in healthy aging. J. Neurosci. 29, 15223–15231.

García-García, I., Michaud, A., Dadar, M., Zeighami, Y., Neseliler, S., Collins, D.L., Evans, A.C., and Dagher, A. (2018). Neuroanatomical differences in obesity: meta-analytic findings and their validation in an independent dataset. Int. J. Obes. 1.

García-García, I., Morys, F., and Dagher, A. (2019). Nucleus accumbens volume is related to obesity measures in an age-dependent fashion. J. Neuroendocrinol.

Gouw, A.A., Seewann, A., Van Der Flier, W.M., Barkhof, F., Rozemuller, A.M., Scheltens, P., and Geurts, J.J.G. (2011). Heterogeneity of small vessel disease: A systematic review of MRI and histopathology correlations. J. Neurol. Neurosurg. Psychiatry 82, 126–135.

Griffanti, L., Zamboni, G., Khan, A., Li, L., Bonifacio, G., Sundaresan, V., Schulz, U.G., Kuker, W., Battaglini, M., Rothwell, P.M., et al. (2016). BIANCA (Brain Intensity AbNormality Classification Algorithm): A new tool for automated segmentation of white matter hyperintensities. Neuroimage 141, 191–205.

Guillemot-Legris, O., and Muccioli, G.G. (2017). Obesity-induced neuroinflammation: beyond the hypothalamus. Trends Neurosci.

Gustafson, D., Lissner, L., Bengtsson, C., Björkelund, C., and Skoog, I. (2004). A 24-year follow-up of body mass index and cerebral atrophy. Neurology 63, 1876–1881.

Hakim, A.M. (2019). Small vessel disease. Front. Neurol. 10.

Hamer, M., and Batty, G.D. (2019). Association of body mass index and waist-to-hip ratio with brain structure. Neurology 92, e594–e600.

Hammond, R.A., and Levine, R. (2010). The economic impact of obesity in the United States. Diabetes. Metab. Syndr. Obes. 3, 285–295.

Han, J.E., Boachie, N., Garcia-Garcia, I., Michaud, A., and Dagher, A. (2018). Neural correlates of dietary self-control in healthy adults: A meta-analysis of functional brain imaging studies. Physiol. Behav. 192, 98–108.

Heneka, M.T., Carson, M.J., Khoury, J. El, Landreth, G.E., Brosseron, F., Feinstein, D.L., Jacobs, A.H., Wyss-Coray, T., Vitorica, J., Ransohoff, R.M., et al. (2015). Neuroinflammation in Alzheimer’s disease. Lancet Neurol. 14, 388–405.

Herrmann, M.J., Tesar, A.K., Beier, J., Berg, M., and Warrings, B. (2019). Grey matter alterations in obesity: A meta-analysis of whole-brain studies. Obes. Rev. 20, 464–471.

Hoaglin, D.C., and Iglewicz, B. (1987). Fine-Tuning Some Resistant Rules for Outlier Labeling. J. Am. Stat. Assoc. 82, 1147–1149.

Hoaglin, D.C., Iglewicz, B., and Tukey, J.W. (1986). Performance of Some Resistant Rules for Outlier Labeling. J. Am. Stat. Assoc. 81, 991–999.

Horstmann, A., Busse, F.P., Mathar, D., Muller, K., Lepsien, J., Schlogl, H., Kabisch, S., Kratzsch, J., Neumann, J., Stumvoll, M., et al. (2011). Obesity-Related Differences between Women and Men in Brain Structure and Goal-Directed Behavior. Front. Hum. Neurosci. 5, 58.

Horstmann, A., Fenske, W.K., and Hankir, M.K. (2015). Argument for a non-linear relationship between severity of human obesity and dopaminergic tone. Obes. Rev. n/a-n/a.

Hsuchou, H., Kastin, A.J., Mishra, P.K., and Pan, W. (2012a). C-reactive protein increases BBB permeability: Implications for obesity and neuroinfammation. Cell. Physiol. Biochem. 30, 1109–1119.

Hsuchou, H., Kastin, A.J., and Pan, W. (2012b). Blood-borne metabolic factors in obesity exacerbate injury-induced gliosis. J. Mol. Neurosci. 47, 267–277.

Hua, P., Pan, X.P., Hu, R., Mo, X.E., Shang, X.Y., and Yang, S.R. (2014). Factors related to executive dysfunction after acute infarct. PLoS One 9.

Hwang, Y.C., Fujimoto, W.Y., Hayashi, T., Kahn, S.E., Leonetti, D.L., and Boyko, E.J. (2016). Increased visceral adipose tissue is an independent predictor for future development of atherogenic dyslipidemia. J. Clin. Endocrinol. Metab. 101, 678–685.

Jenkinson, M., Beckmann, C.F., Behrens, T.E.J., Woolrich, M.W., and Smith, S.M. (2012). FSL. Neuroimage 62, 782–790.

Kakoschke, N., Lorenzetti, V., Caeyenberghs, K., and Verdejo-García, A. (2019). Impulsivity and body fat accumulation are linked to cortical and subcortical brain volumes among adolescents and adults. Sci. Rep. 9, 1–11.

Kharabian Masouleh, S., Arélin, K., Horstmann, A., Lampe, L., Kipping, J.A., Luck, T., Riedel-Heller, S.G., Schroeter, M.L., Stumvoll, M., Villringer, A., et al. (2016). Higher body mass index in older adults is associated with lower gray matter volume: Implications for memory performance. Neurobiol. Aging 40, 1–10.

Kivipelto, M., Ngandu, T., Fratiglioni, L., Viitanen, M., Kåreholt, I., Winblad, B., Helkala, E.L., Tuomilehto, J., Soininen, H., and Nissinen, A. (2005). Obesity and vascular risk factors at midlife and the risk of dementia and Alzheimer disease. Arch. Neurol. 62, 1556–1560.

Klein, A., and Tourville, J. (2012). 101 labeled brain images and a consistent human cortical labeling protocol. Front. Neurosci. 6.

Kuo, H.K., Chen, C.Y., Liu, H.M., Yen, C.J., Chang, K.J., Chang, C.C., Yu, Y.H., Lin, L.Y., and Hwang, J.J. (2010). Metabolic risks, white matter hyperintensities, and arterial stiffness in high-functioning healthy adults. Int. J. Cardiol. 143, 184–191.

Lam, L.C.W., Chan, W.C., Leung, T., Fung, A.W.T., and Leung, E.M.F. (2015). Would older adults with mild cognitive impairment adhere to and benefit from a structured lifestyle activity intervention to enhance cognition?: A cluster randomized controlled trial. PLoS One 10.

Lambert, C., Sam Narean, J., Benjamin, P., Zeestraten, E., Barrick, T.R., and Markus, H.S. (2015). Characterising the grey matter correlates of leukoaraiosis in cerebral small vessel disease. NeuroImage Clin. 9, 194–205.

Lampe, L., Zhang, R., Beyer, F., Huhn, S., Kharabian□Masouleh, S., Preusser, S., Bazin, P., Schroeter, M.L., Villringer, A., and Witte, A.V. (2018). Visceral Obesity Relates to Deep White Matter Hyperintensities via Inflammation. Ann. Neurol. 85, ana.25396.

Lee, K.S., Lee, Y., Back, J.H., Son, S.J., Choi, S.H., Chung, Y.K., Lim, K.Y., Noh, J.S., Koh, S.H., Oh, B.H., et al. (2014). Effects of a multidomain lifestyle modification on cognitive function in older adults: An eighteen-month community-based cluster randomized controlled trial. Psychother. Psychosom. 83, 270–278.

de Leeuw, F.-E., de Groot, J.C., Oudkerk, M., Witteman, J.C.M., Hofman, A., van Gijn, J., and Breteler, M.M.B. (2002). Hypertension and cerebral white matter lesions in a prospective cohort study. Brain 125, 765–772.

Madden, D.J., Bennett, I.J., Burzynska, A., Potter, G.G., Chen, N. kuei, and Song, A.W. (2012). Diffusion tensor imaging of cerebral white matter integrity in cognitive aging. Biochim. Biophys. Acta - Mol. Basis Dis. 1822, 386–400.

Maldonado-Ruiz, R., Montalvo-Martínez, L., Fuentes-Mera, L., and Camacho, A. (2017). Microglia activation due to obesity programs metabolic failure leading to type two diabetes. Nutr. Diabetes 7, e254–e254.

Mathieu, P., Poirier, P., Pibarot, P., Lemieux, I., and Després, J.-P. (2009). Visceral obesity: the link among inflammation, hypertension, and cardiovascular disease. Hypertens. (Dallas, Tex. 1979) 53, 577–584.

Medic, N., Ziauddeen, H., Ersche, K.D., Farooqi, I.S., Bullmore, E.T., Nathan, P.J., Ronan, L., and Fletcher, P.C. (2016). Increased body mass index is associated with specific regional alterations in brain structure. Int. J. Obes. 40, 1177–1182.

Mehl, N., Morys, F., Villringer, A., and Horstmann, A. (2019). Unhealthy yet avoidable — how cognitive bias modification alters behavioral and brain responses. Nutrients 11.

Miller, A.A., and Spencer, S.J. (2014). Obesity and neuroinflammation: A pathway to cognitive impairment. Brain. Behav. Immun. 42, 10–21.

Miller, K.L., Alfaro-Almagro, F., Bangerter, N.K., Thomas, D.L., Yacoub, E., Xu, J., Bartsch, A.J., Jbabdi, S., Sotiropoulos, S.N., Andersson, J.L.R., et al. (2016). Multimodal population brain imaging in the UK Biobank prospective epidemiological study. Nat. Neurosci. 19, 1523– 1536.

Monteiro, R., and Azevedo, I. (2010). Chronic inflammation in obesity and the metabolic syndrome. Mediators Inflamm. 2010.

Mori, S., Wakana, S., Nagae-Poetscher, L., and van Zijl, P. (2005). MRI atlas of human white matter (Amsterdam: Elsevier B.V.).

Neseliler, S., Vainik, U., Dadar, M., Yau, Y.H.C., Garcia-Garcia, I., Scala, S.G., Zeighami, Y., Collins, D.L., and Dagher, A. (2018). Neural and behavioral endophenotypes of obesity. BioRxiv 348821.

Ngandu, T., Lehtisalo, J., Solomon, A., Levälahti, E., Ahtiluoto, S., Antikainen, R., Bäckman, L., Hänninen, T., Jula, A., Laatikainen, T., et al. (2015). A 2 year multidomain intervention of diet, exercise, cognitive training, and vascular risk monitoring versus control to prevent cognitive decline in at-risk elderly people (FINGER): A randomised controlled trial. Lancet 385, 2255–2263.

Nguyen, J.C.D., Killcross, A.S., and Jenkins, T.A. (2014). Obesity and cognitive decline: Role of inflammation and vascular changes. Front. Neurosci. 8.

Noble, R.E. (2001). Waist-to-hip ratio versus BMI as predictors of cardiac risk in obese adult women. West. J. Med. 174, 240-a-241.

Pannacciulli, N., Del Parigi, A., Chen, K., Le, D.S.N.T., Reiman, E.M., and Tataranni, P.A. (2006). Brain abnormalities in human obesity: A voxel-based morphometric study. Neuroimage 31, 1419–1425.

Pantoni, L. (2010). Cerebral small vessel disease: from pathogenesis and clinical characteristics to therapeutic challenges. Lancet Neurol. 9, 689–701.

Pareek, V., Rallabandi, V.S., and Roy, P.K. (2018). A Correlational Study between Microstructural White Matter Properties and Macrostructural Gray Matter Volume Across Normal Ageing: Conjoint DTI and VBM Analysis. Magn. Reson. Insights 11, 1178623X1879992.

Patenaude, B., Smith, S.M., Kennedy, D.N., and Jenkinson, M. (2011). A Bayesian model of shape and appearance for subcortical brain segmentation. Neuroimage 56, 907–922.

Rapuano, K.M., Zieselman, A.L., Kelley, W.M., Sargent, J.D., Heatherton, T.F., and Gilbert-Diamond, D. (2017). Genetic risk for obesity predicts nucleus accumbens size and responsivity to real-world food cues. Proc. Natl. Acad. Sci. 114, 160–165.

Ronan, L., Alexander-Bloch, A.F., Wagstyl, K., Farooqi, S., Brayne, C., Tyler, L.K., and Fletcher, P.C. (2016). Obesity associated with increased brain age from midlife. Neurobiol. Aging 47, 63–70.

Rosseel, Y. (2012). lavaan: An R Package for Structural Equation Modelinge human forearm during rythmic exercise. J. Stat. Softw. 48, 1–36.

Samara, A., Murphy, T., Strain, J., Rutlin, J., Sun, P., Neyman, O., Sreevalsan, N., Shimony, J.S., Ances, B.M., Song, S.K., et al. (2020). Neuroinflammation and White Matter Alterations in Obesity Assessed by Diffusion Basis Spectrum Imaging. Front. Hum. Neurosci. 13, 464.

Sen, P.N., and Basser, P.J. (2005). A model for diffusion in white matter in the brain. Biophys. J. 89, 2927–2938.

Shaw, M.E., Sachdev, P.S., Abhayaratna, W., Anstey, K.J., and Cherbuin, N. (2018). Body mass index is associated with cortical thinning with different patterns in mid- and late-life. Int. J. Obes. 42, 455–461.

Singh, V., Chertkow, H., Lerch, J.P., Evans, A.C., Dorr, A.E., and Kabani, N.J. (2006). Spatial patterns of cortical thinning in mild cognitive impairment and Alzheimer’s disease. Brain 129, 2885–2893.

Smith, C.E., and Cribbie, R.A. (2013). Multiplicity Control in Structural Equation Modeling: Incorporating Parameter Dependencies. Struct. Equ. Model. A Multidiscip. J. 20, 79–85.

Soares, J.M., Marques, P., Alves, V., and Sousa, N. (2013). A hitchhiker’s guide to diffusion tensor imaging. Front. Neurosci. 7.

Song, S.K., Sun, S.W., Ramsbottom, M.J., Chang, C., Russell, J., and Cross, A.H. (2002). Dysmyelination revealed through MRI as increased radial (but unchanged axial) diffusion of water. Neuroimage 17, 1429–1436.

Sproston, N.R., and Ashworth, J.J. (2018). Role of C-reactive protein at sites of inflammation and infection. Front. Immunol. 9.

Spyridaki, E.C., Simos, P., Avgoustinaki, P.D., Dermitzaki, E., Venihaki, M., Bardos, A.N., and Margioris, A.N. (2014). The association between obesity and fluid intelligence impairment is mediated by chronic low-grade inflammation. Br. J. Nutr. 112, 1724–1734.

Streit, W.J., Walter, S.A., and Pennell, N.A. (1999). Reactive microgliosis. Prog. Neurobiol. 57, 563–581.

Sudlow, C., Gallacher, J., Allen, N., Beral, V., Burton, P., Danesh, J., Downey, P., Elliott, P., Green, J., Landray, M., et al. (2015). UK Biobank: An Open Access Resource for Identifying the Causes of a Wide Range of Complex Diseases of Middle and Old Age. PLOS Med. 12, e1001779.

Swardfager, W., Yu, D., Ramirez, J., Cogo-Moreira, H., Szilagyi, G., Holmes, M.F., Scott, C.J.M., Scola, G., Chan, P.C., Chen, J., et al. (2017). Peripheral inflammatory markers indicate microstructural damage within periventricular white matter hyperintensities in Alzheimer’s disease: A preliminary report. Alzheimer’s Dement. Diagnosis, Assess. Dis. Monit. 7, 56–60.

Taki, Y., Kinomura, S., Sato, K., Inoue, K., Goto, R., Okada, K., Uchida, S., Kawashima, R., and Fukuda, H. (2008). Relationship Between Body Mass Index and Gray Matter Volume in 1,428 Healthy Individuals. Obesity 16, 119–124.

Tamura, Y., and Araki, A. (2015). Diabetes mellitus and white matter hyperintensity. Geriatr. Gerontol. Int. 15, 34–42.

Tingley, D., Yamamoto, T., Hirose, K., Keele, L., and Imai, K. (2014). Mediation: R package for causal mediation analysis. J. Stat. Softw. 59, 1–38.

Tukey, J.W. (1977). Exploratory Data Analysis (Reading, Mass: Pearson).

Tuladhar, A.M., Reid, A.T., Shumskaya, E., De Laat, K.F., Van Norden, A.G.W., Van Dijk, E.J., Norris, D.G., and De Leeuw, F.E. (2015). Relationship between white matter hyperintensities, cortical thickness, and cognition. Stroke 46, 425–432.

Vainik, U., Baker, T.E., Dadar, M., Zeighami, Y., Michaud, A., Zhang, Y., Alanis, J.C.G., Misic, B., Collins, D.L., and Dagher, A. (2018). Neurobehavioral correlates of obesity are largely heritable. Proc. Natl. Acad. Sci. 201718206.

Veit, R., Kullmann, S., Heni, M., Machann, J., Häring, H.U., Fritsche, A., and Preissl, H. (2014). Reduced cortical thickness associated with visceral fat and BMI. NeuroImage Clin. 6, 307–311.

Visser, M., Bouter, L.M., McQuillan, G.M., Wener, M.H., and Harris, T.B. (2001). Low-grade systemic inflammation in overweight children. Pediatrics 107, E13.

Volkow, N.D., Wang, G.-J., Fowler, J.S., and Telang, F. (2008). Overlapping Neuronal Circuits in Addiction and Obesity: Evidence of Systems Pathology. Philos. Trans. Biol. Sci. 363, 3191–3200.

Wakana, S., Caprihan, A., Panzenboeck, M.M., Fallon, J.H., Perry, M., Gollub, R.L., Hua, K., Zhang, J., Jiang, H., Dubey, P., et al. (2007). Reproducibility of quantitative tractography methods applied to cerebral white matter. Neuroimage 36, 630–644.

Walker, K.A., Power, M.C., Hoogeveen, R.C., Folsom, A.R., Ballantyne, C.M., Knopman, D.S., Windham, B.G., Selvin, E., Jack, C.R., and Gottesman, R.F. (2017). Midlife systemic inflammation, late-life white matter integrity, and cerebral small vessel disease the atherosclerosis risk in communities study. Stroke 48, 3196–3202.

Wang, H., Wen, B., Cheng, J., and Li, H. (2017). Brain structural differences between normal and obese adults and their links with lack of perseverance, negative urgency, and sensation seeking. Sci. Rep. 7.

Ward, M.A., Carlsson, C.M., Trivedi, M.A., Sager, M.A., and Johnson, S.C. (2005). The effect of body mass index on global brain volume in middle-aged adults: A cross sectional study. BMC Neurol. 5, 23.

Wardlaw, J.M., Valdés Hernández, M.C., and Muñoz-Maniega, S. (2015). What are white matter hyperintensities made of? Relevance to vascular cognitive impairment. J. Am. Heart Assoc. 4, 001140.

Wardlaw, J.M., Makin, S.J., Valdés Hernández, M.C., Armitage, P.A., Heye, A.K., Chappell, F.M., Muñoz-Maniega, S., Sakka, E., Shuler, K., Dennis, M.S., et al. (2017). Blood-brain barrier failure as a core mechanism in cerebral small vessel disease and dementia: evidence from a cohort study. Alzheimer’s Dement. 13, 634–643.

Whitmer, R., Gunderson, E., Quesenberry, C., Zhou, J., and Yaffe, K. (2007). Body Mass Index in Midlife and Risk of Alzheimer Disease and Vascular Dementia. Curr. Alzheimer Res. 4, 103–109.

Yaffe, K., Kanaya, A., Lindquist, K., Simonsick, E.M., Harris, T., Shorr, R.I., Tylavsky, F.A., and Newman, A.B. (2004). The metabolic syndrome, inflammation, and risk of cognitive decline. J. Am. Med. Assoc. 292, 2237–2242.

Zhang, R., Beyer, F., Lampe, L., Luck, T., Riedel-Heller, S.G., Loeffler, M., Schroeter, M.L., Stumvoll, M., Villringer, A., and Witte, A.V. (2018). White matter microstructural variability mediates the relation between obesity and cognition in healthy adults. Neuroimage 172, 239–249.

Zhao, Y., Ke, Z., He, W., and Cai, Z. (2019). Volume of white matter hyperintensities increases with blood pressure in patients with hypertension. J. Int. Med. Res. 47, 3681–3689.

Zhou, J., and Qin, G. (2012). Adipocyte dysfunction and hypertension. Am. J. Cardiovasc. Dis. 2, 143–149.

